# General structure-free energy relationships of hERG blocker binding under native cellular conditions

**DOI:** 10.1101/2021.10.07.463585

**Authors:** Hongbin Wan, Kristina Spiru, Sarah Williams, Robert A. Pearlstein

## Abstract

We proposed previously that aqueous non-covalent barriers arise from solute-induced perturbation of the H-bond network of solvating water (“the solvation field”) relative to bulk solvent, where the association barrier equates to enthalpic losses incurred from incomplete replacement of the H-bonds of expelled H-bond enriched solvation by inter-partner H-bonds, and the dissociation barrier equates to enthalpic + entropic losses incurred during dissociation-induced resolvation of H-bond depleted positions of the free partners (where dynamic occupancy is powered largely by the expulsion of such solvation to bulk solvent during association). We went on to analyze blockade of the human ether-a-go-go-related gene potassium channel (hERG) based on these principles, the results of which suggest that blockers: 1) project a single rod-shaped R-group (denoted as “BP”) into the pore at a rate proportional to the desolvation cost of BP, with the largely solvated remainder (denoted as “BC”) occupying the cytoplasmic “antechamber” of hERG; and 2) undergo second-order entry to the antechamber, followed by <u>first-order</u> association of BP to the pore. In this work, we used WATMD to qualitatively survey the solvation fields of the pore and a representative set of 16 blockers sampled from the Redfern dataset of marketed drugs spanning a range of pro-arrhythmicity. We show that the highly non-polar <u>pore</u> is solvated principally by H-bond depleted and bulk-like water (incurring zero desolvation cost), whereas <u>blocker BP moieties</u> are solvated by variable combinations of H-bond enriched and depleted water. With a few explainable exceptions, the <u>blocker</u> solvation fields (and implied desolvation/resolvation costs) are qualitatively well-correlated with both blocker potency and Redfern safety classification.

## Introduction

Despite many years of intensive investigation into the possible causes of, and remedies for, inadvertent blockade of the hERG potassium channel by chemically diverse low molecular weight (LMW) hits, leads, preclinical/clinical candidates, and drugs, hERG blockade remains one of the many unsolved liabilities that are typically addressed via black box trial-and-error workflows. However, this approach is hampered by the convoluted nature of target/off-target potency, solubility, permeability, and pharmacokinetics (PK), in which modulation of one property or behavior can positively or negatively affect one or more of the others. Lead optimization often culminates in residual hERG activity at the clinical candidate stage, resulting in potential no-go decisions or mandated clinical thorough QT (TQT) studies, depending on the benefit/risk ratio.

We previously proposed that the canonical hERG blocker motif consists of a chemically diverse Y-shaped topology similar to that proposed by Cavalli et al. [1]:

1. The <u>stem</u> of the Y consists of a quasi-rod-shaped moiety (denoted as “BP”) that transiently projects into the ion conduction pathway or pore of hERG (denoted as “P”) [2]. Basic groups that are prevalent among hERG blockers always reside somewhere within this moiety.
2. The V-shaped <u>cap</u> of the Y (denoted as “BC”) resides within the large cytoplasmic “antechamber” (denoted as “C”) adjoining the pore entrance, which is lined by the C-linker and cyclic nucleotide binding homology (CNBH) domains. BP and BC straddle between P and C, respectively

The lack of significant progress toward the development of reliable strategies for hERG avoidance and mitigation may be attributed to:

1. Poor <u>general</u> understanding of non-covalent binding between cognate partners, including drugs and targets/off-targets (described below).
2. Consideration of hERG blockade as a <u>typical</u> binding process, when in fact, it is highly atypical due to:

a. <u>The absence of native binding function of the ion conduction pathway</u>, which serves as the binding site for all known blockers. We attribute native binding function, in general, to complementarity between steric size/shape and the solvation properties of cognate binding partners described below [3–5].
b. <u>Two-step binding</u>, consisting of:

i. The capture of a single solvated blocker copy within C. The on-rate is described by k_c_ · [free antechamber] · [free blocker] (typical <u>second-order</u> binding, in which k_on_ is capped at the 10^9^ M^-1^ s^-1^ diffusion limit), where k_c_ denotes the blocker-antechamber association rate constant. The blocker-bound antechamber concentration builds and decays with the free cytoplasmic concentration, which in turn, builds and decays with cardiomyocyte uptake and plasma clearance, respectively.
ii. Transient projection of BP into the open pore [2]. In this step, the on-rate is described by k_b_ · [antechamber-bound blocker] (atypical <u>first-order</u> binding), where k_b_ denotes the BP-P association rate constant, and the BP off-rate (to the antechamber) is described by k_-b_ · [bound blocker], where k_-b_ is the BP-P dissociation rate constant. Many BP-P association/dissociation cycles may occur for each captured blocker prior to dissociation from C.
3. The lack of differentiation between the safety profiles of trappable blockers that remain bound in closed channels (where BP and BC reside on opposite sides of the closed activation gate) and non-trappable blockers that are expelled during channel closing, together with poor understanding of structure-trappability relationships [2,6,7]. Fractional occupancy differs kinetically among the two blocker sub-types as follows:

a. <u>Non-trappables</u>: occupancy builds and decays during each channel gating cycle, the peak magnitude of which occurs at the intracellular C_max_ (the peak exposure during each dosing interval), which in turn builds and decays during the distribution and clearance phases of the PK curve, respectively. The highest maximum occupancy of non-trappable blockers corresponds to k_b_ ≈ the rate of channel opening and k_-b_ ≈ the rate of channel closing (where k_-b_ is usurped by the rate of channel closing).
b. <u>Trappables</u>: occupancy accumulates to the maximum over multiple channel gating cycles, given approximately by the Hill equation (free C_max_/(free C_max_ + IC_50_), in which the exponents are assumed = 1) as the intracellular C_max_ builds to n · IC_50_, where n is the occupancy multiplier (e.g., n = 1 equates to 50% occupancy, n = 19 equates to 95% occupancy, etc.). Occupancy decays during the clearance phase of the PK curve (noting that k_-b_ is <u>not</u> usurped by the rate of channel closing).
4. <u>Measurement of hERG blockade under static equilibrium conditions</u> in the status quo hERG assays, and the use of such data for generating hERG structure-activity relationship (SAR) models when, in reality, native binding between hERG and <u>non-trappable</u> blockers operates in the highly non-equilibrium regime. In our previous work, we simulated virtual hERG blockade (non-structurally) in the context of the cardiac AP using a version of the O’Hara-Rudy model of undiseased human ventricular cardiomyocytes [8,9] that we modified [10,11]. We showed that hERG occupancy by non-trappable blockers depends far more on fast k_b_ than slow k_-b_ due to fast buildup of the open state of the pore [10,11]. Under native conditions, typical drug-target systems operate on far longer timescales than the ∼350 ms open channel time window of hERG, commensurate with significantly slower requirements for k_on_ and k_off_.
5. <u>Key deficiencies in the “Redfern hERG safety index”</u> (hERG IC_50_/free C_max_ > 30) that was derived from PK, IC_50_, and adverse clinical event data reported for 52 marketed drugs. In our previous work, we showed that the Redfern safety index (SI) equates to near zero safe fractional hERG occupancy at the therapeutic free C_max_, allowing for unintended exposure escalation due to overdose or drug-drug interactions (DDIs) (noting that all reported hERG blockade-induced arrhythmia cases in humans involved aberrant blood levels <u>in excess of</u> the therapeutic free C_max_).
6. Mutual desolvation of P and BP during association depends on the existence of a water-accessible pathway between P and bulk-solvent. Since no obvious pathway exists at the extracellular end of P, displaced water is necessarily expelled from the free volume of the blocker-bound pore through the pore entrance, thereby precluding total occlusion of P by BP in fully buried blocker-bound states proposed elsewhere (e.g., [10,12,13]).

The development of improved mitigation and safety assessment strategies, compared with the status quo black box trial-and-error approaches, depends on achieving a deeper understanding of the true mechanisms of hERG blockade under native cellular conditions.

### Binding free energy is contributed principally by solvation, and the implications thereof for hERG blockade

Mounting evidence suggests that non-covalent intra- and intermolecular rearrangments are powered principally by favorable and unfavorable free energy stored in the hydrogen bonds (H-bonds) of water solvating the external surfaces of LMW and high molecular weight (HMW) solutes, as well as within the internal cavities of HMW solutes (rather than in interatomic contacts) [3–5,14–17]. In our previous work, we postulated that water H-bond networks behave as dynamic fields, in which:

1. The number/strength (enthalpy) and ordering (entropy) of water-water and water-solute H-bonds vary at each position of the field <u>relative to those of bulk solvent</u>.
2. The free energy of each solvating water is enthalpically enriched, depleted, or neutral, depending on the local solute surface composition and topology (noting that all solvating water is entropically depleted, resulting in entropy-enthalpy compensation).
3. The rates of water exchanges between the field and bulk solvent (which we refer to as “solvation dynamics”) vary among enthalpically enriched, depleted, and bulk-like neutral positions, as follows:

a. H-bond depleted solvation undergoes slow water exchanges from bulk solvent to the solvation field and fast water exchanges from the solvation field to bulk solvent.
b. H-bond enriched solvation undergoes fast water exchanges from bulk solvent to the solvation field and slow water exchanges from the solvation to bulk solvent.
c. Bulk-like (energetically neutral) solvation undergoes equal rates of exchange to/from bulk solvent.

We further postulated that:

1. The <u>entry/association</u> barrier to non-covalent intra-/intermolecular states consists principally of the total free energy cost of <u>desolvating</u> H-bond <u>enriched</u> solvation from the rearrangement or binding interface (to which k_in_ and k_on_ are proportional), and the canonical <u>exit/dissociation</u> barrier consists principally of the free energy cost of <u>resolvating</u> H-bond <u>depleted</u> solvation positions that were vacated during entry/association (to which k_out_ and k_off_ are proportional) [3–5,14,15,17,18]. These costs consist of the following:

a. The desolvation cost depends on the degree to which the H-bonds of H-bond enriched water solvating the pre-bound partners are mutually replaced by polar groups in the bound complex (noting that the free energy gains from such replacements are typically ≤ the free energy lost from desolvation, resulting in a zero sum game at best). The greater the degree of H-bond enrichment, the more stringent are the H-bond replacement requirements. H-bond enriched solvating water thus serves as the “gatekeeper” for entry/association to rearrangement or binding interfaces and the basis for specificity and solubility (noting that solubility is directly proportional to the solvation free energy, which in turn, is equal to the desolvation cost).
b. The resolvation cost depends on the degree to which H-bond depleted solvation in the pre-bound state is replaced by non-polar or weakly polar groups in the bound state. The desolvation gain from the expulsion of H-bond depleted solvation is the principal driving force for non-covalent intra- and intermolecular rearrangements, wherein the greater the degree of H-bond depletion, the higher the resolvation cost per unit area of the exited or dissociated partners, respectively. We refer to the spatial arrangement and free energy distribution of the solvating water within functional rearrangement and binding interfaces as “solvophores”. It follows that solvophores are absent in non-native binding sites, including the ion conduction pathway of hERG.
2. Solubility and logP depend on the balance and surface distribution of H-bond enriched and depleted solvation (neglecting the cost of dissolution), and are therefore vectorial rather than scalar quantities (e.g., logP, solubility). High solubility translates to a high <u>desolvation</u> cost, resulting in slower k_in_ or k_on_ to rearrangement or binding interfaces and slow k_in_ to membrane surfaces. Low solubility translates to a high <u>resolvation</u> cost, resulting in slow k_out_ or k_off_.
3. Permeability likewise depends on the desolvation and resolvation costs of both the permeants and membrane surfaces. The k_in_ for membrane surface penetration depends on the desolvation costs of both permeants and membrane surfaces (noting that <u>blockers</u> necessarily enter hERG from the intracellular opening).

Based on our earlier work with WaterMap, we predicted that P is solvated predominantly by bulk-like and H-bond depleted solvation localized around the side chains of Phe656 and Tyr652 [10], which is consistent with:

1. The largely non-polar composition of the lumen.
2. The attenuating effect of ordered, H-bond enriched solvation on the negative electrostatic potential within the ion conduction pathway [10].
3. The high promiscuity of the channel in the absence of H-bond enriched “gatekeeper” solvation, relegating the association barrier principally to steric size/shape complementarity between P and BP and the desolvation cost of <u>BP</u> [3] (noting that binding is largely non-specific in the absence of H-bond enriched “gatekeeper” solvation).

In this work, we used WATMD (described briefly in Materials and methods and fully elsewhere [4,16]) to characterize the solvation dynamics of P, C, BP, and BC among a representative subset of hERG-blocking drugs exhibiting a wide range of potencies and pro-arrhythmicities. We show that:

1. BP is solvated by both H-bond enriched and depleted solvation, as reflected in the presence of both high and ultra-low occupancy voxels (denoted as HOVs and ULOVs, respectively).
2. P is solvated almost exclusively by H-bond depleted solvation, as reflected in the nearly complete absence of ULOVs.
3. Blockers are only partially desolvated during transient pore association [2]. Therefore, knowing where to increase the desolvation cost on blocker surfaces is essential for “surgical” hERG mitigation.
4. k_b_ and k_-b_ governing blocker potency are qualitatively proportional to the degree of H-bond enriched solvation (HOVs) of BP and depleted solvation (ULOVs) of P and BP, given that:

a. The desolvation cost of P is nearly zero in the absence of H-bond enriched solvation, and therefore contributes little to the total BP + P desolvation cost.
b. The desolvation cost of BP is maximal in the absence of polar groups lining P needed to replace the H-bonds of the H-bond enriched solvation of BP.

## Materials and methods

Molecular dynamics (MD) simulations are used extensively for predicting the intra- and intermolecular structural rearrangements of proteins and other biomolecules [19–22]. However, our simulations are focused solely on water exchanges between solvation and bulk solvent (which we refer to as “solvation dynamics (SD) simulations”). We used WATMD (fully described in reference 4) to calculate the solvation fields of a set of marketed/withdrawn drugs exhibiting known hERG activity and pro-arrhythmicities selected from the dataset compiled by Redfern et al. (Table 1) [23]. Briefly, WATMD counts the number of visits of water oxygen (O) and hydrogen (H) atoms to each of a set of 1 Å^3^ cells (voxels) within a three-dimensional lattice fully surrounding a LMW or HMW solute of interest during the last 10 ns of a 100 ns MD simulation. The counts per voxel are always distributed in a Gaussian-like fashion around the mean H and O counts corresponding to bulk-like solvation (normalized for the 2 H/O ratio). Voxels exhibiting ultra-low counts (ULOVs) and ultra-high counts (HOVs) fall within the left and right tails of the distribution [4]. ULOVs and HOVs are represented as spheres, the radii of which are proportional to the counts, and the colors of which are assigned as follows:

1. Bright red = 100% O visits.
2. Varying shades of pink = mixture of O and H visits, tipped toward O.
3. White = no preference for O or H. All ULOVs are colored white due to the typically non-polar environment of these voxels. Alternatively, white HOVs may be indicative of mixed donor/acceptor voxel environments or water molecules that are trapped within non-polar cavities (depending on the context).
4. Varying shades of purple = mixture of O and H visits, tipped toward H.
5. Bright blue = 100% H visits.

**Table 1.** A subset of the hERG blockers and data compiled by Redfern et al. selected for our study. The data consists of the adverse event Class assigned by those authors (1 = antiarrhythmic drugs that were also to be pro-arrhythmic; 2 = drugs that were withdrawn due to high arrhythmic risk/benefit ratio; 3 = drugs with numerous reported cases of arrhythmia; 4 = drugs with isolated reports of arrhythmia; 5 = drugs with no reported cases of arrhythmia), minimum and maximum reported hERG IC_50_, and minimum and maximum reported plasma C_max_. Trappable and non-trappable blockers reported by Stork et al. [6] and Windisch et al. [7] are denoted by * and #, respectively.

| **Blocker** | **Class** | **hERG IC<sub>50</sub>**<br>**range (μM)** |  | **Upper C<sub>max</sub>**<br>**range (μM)** |  | **Min hERG**<br>**IC<sub>50</sub>/max C<sub>max</sub>** |
| --- | --- | --- | --- | --- | --- | --- |
| quinidine | 1 | 0.3 | 1.0 | 0.92 | 3.2 | **0.09** |
| ibutilide | 1 | 0.01 | 0.02 | 7e-4 | 0.14 | **0.07** |
| almokalant | 1 | 0.05 | - | 0.07 | 0.15 | **0.33** |
| sertindole | 2 | 0.014 | 0.062 | 2e-5 | 1.6e-3 | 8.75 |
| terfenadine* | 2 | 0.02 | 0.2 | 0.1 | 0.29 | **0.07** |
| cisapride# | 2 | 0.002 | 0.045 | 2.6e-3 | 4.9e-3 | **0.41** |
| terodiline | 2 | 0.004 | 0.7 | 8e-3 | 0.012 | **0.33** |
| thioridazine | 3 | 0.033 | 1.25 | 0.21 | 0.98 | **0.03** |
| pimozide | 3 | 0.015 | 0.055 | 9e-3 | 0.043 | **0.35** |
| flecainide | 3 | 3.91 | - | 0.38 | 0.75 | 5.2 |
| fexofenadine | 4 | 5 | 23 | 0.35 | - | 14.2 |
| imipramine | 4 | 3.4 | - | 0.035 | 0.11 | 30.9 |
| propafenone* | 4 | 0.44 | - | 0.026 | 0.24 | 1.8 |
| desipramine | 4 | 1.39 | - | 0.027 | 0.11 | 12.6 |
| ebastine | 5 | 0.3 | - | 0.04 | 0.05 | 6 |
| verapamil | 5 | 0.14 | 0.83 | 0.025 | 0.08 | 1.75 |

The LMW SD protocol differs from the HMW protocol described in reference 4 in that LMW structures are fully <u>restrained</u> during the simulations (which would otherwise distribute over a large number of non-native conformations in proportion to their force-field-calculated energies), whereas HMW structures are fully <u>unrestrained</u> (self-limited to high frequency rearrangements among the side chains and loops). All blocker structures were generated in their charged forms using the Build Tool of Maestro release 2021-2 (Schrodinger, LLC), and minimized using the default minimization protocol. The structures were overlaid manually on our previously published qualitative blocker overlay model [2] using the manual superposition tool of Maestro. Each structure was then simulated using AMBER 20 PMEMD CUDA (GAFF and ff99sb force-fields) for 100 ns in a box of explicit TIP3P water molecules, and the last 10 ns of each trajectory (40,000 frames) was processed into voxel counts via WATMD, and visualized as spheres using PyMol 2.4.1 (Schrodinger, LLC). We tested the conformational sensitivity of our results by calculating the solvation fields for mildly modified terfenadine and fexofenadine conformations relative to those in the overlay model (noting that significant variation of the solvation fields among highly dissimilar conformations is expected).

The open hERG cryo-EM structure (PDB code = 5VA1 [24]) was prepared using the default settings of the PPrep tool in Maestro. The starting POPC phospholipid membrane-bound structure was taken from reference 13 and subjected to a fully unrestrained 100 ns SD simulation in a box of explicit TIP3P water molecules using AMBER 20 PMEMD CUDA (ff14sb and Lipid14 force-fields). The resulting trajectory was processed into voxel counts as described above and in reference 4). The charged form of astemizole was docked in the astemizole-bound structure (PDB code = 7CN1 [12]) using the Glide XP tool in Maestro (noting that astemizole was omitted from this structure for unexplained reasons). We qualitatively compared the binding modes of quinidine (PDB code = 6LQA [25]) and flecainide (PDB code = 6UZ0 [26]) in Na_v_1.5 with that of astemizole and our binding model.

## Results

We postulated previously that pore occupancy by non-trappable blockers builds and decays transiently during each action potential (AP) cycle, whereas that of trappable blockers accumulates across multiple APs [10,11]. Blocker occupancy is governed by solubility and permeability (underlying the free intracellular blocker concentration), pKa, and the desovlation/resolvation costs of the BP and BC moieties (noting that k_-b_ < the rate of channel closing has no impact on potency under native cellular conditions). Surgical hERG mitigation is often constrained by the typically narrow separation between these properties among therapeutic targets versus hERG and other off-target occupancies, all of which stem directly from (or in the case of pKa are modulated by) the solvation field. Here, we have used WATMD to qualitatively explore the desolvation costs of the blockers listed in Table 1. We postulate that blockers project their BP moieties into P at rates governed by the full and partial desolvation costs of BP and BC, respectively, as reflected qualitatively in the sizes of the HOVs surrounding those moieties. We overlaid the blockers as described in Materials and methods and reference 2 (Figure 1), which we then used to assign the BP and BC moieties and compare the calculated solvation fields across the dataset. The objectives of this work include:

1. Testing our straddled BP-in/BC-out blocker binding geometry hypothesis. If correct, the larger HOVs should be concentrated around the BC moiety of each blocker, which is predicted to remain bound within the well-solvated antechamber (possibly undergoing partial desolvation in the peri-entrance region in the BP-in state [2]).
2. Assessing the relationship between HOV position/size/number and blocker potency. Fewer or smaller HOVs should occur on the BP moieties of Classes 1-3 compared with Class 4-5 drugs.

**Figure 1.**
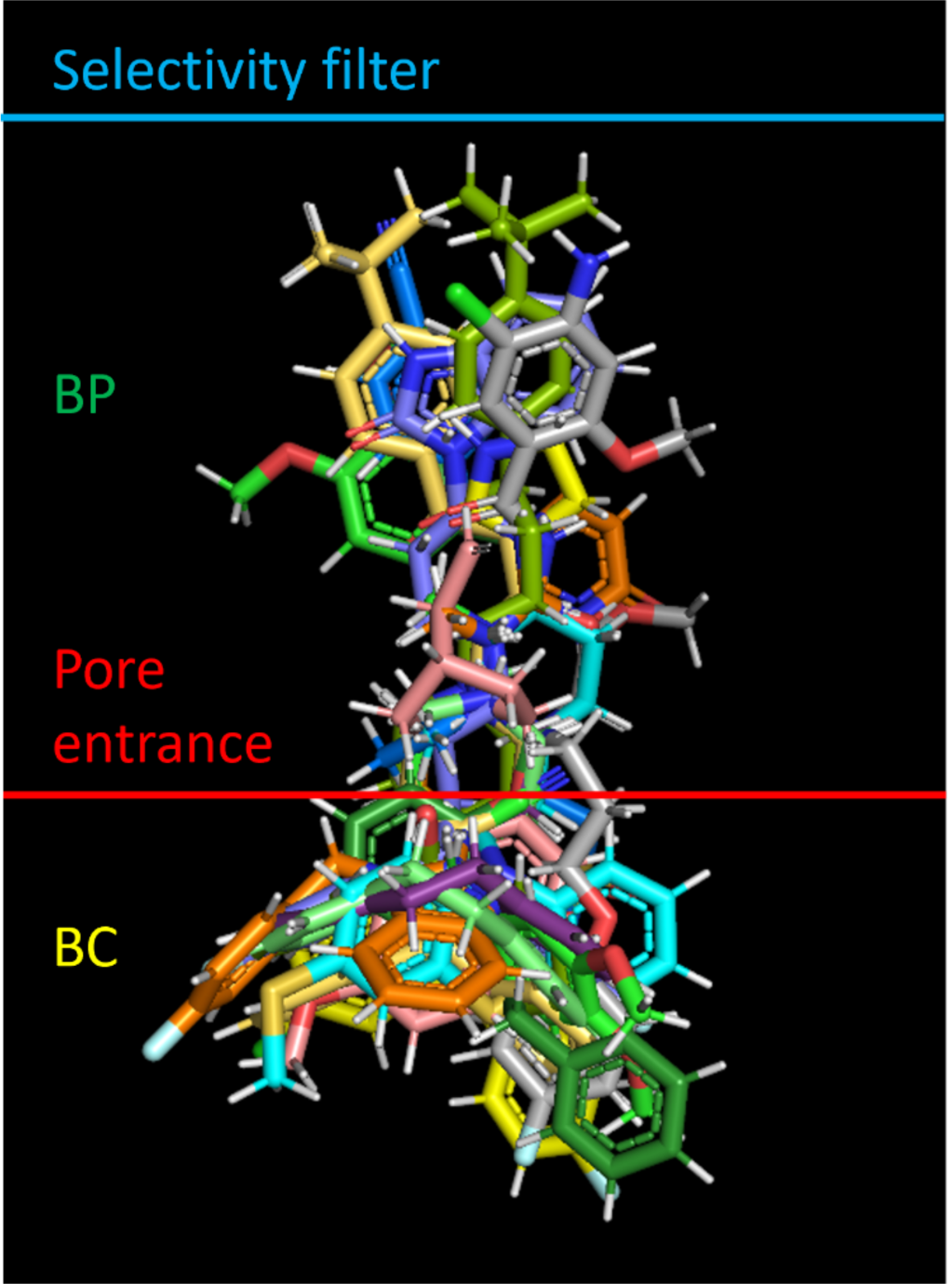
Qualitative manual overlay of the blockers listed in Table 1, as described in Materials and methods and reference 2 (keeping in mind that the distribution of possible bound conformational states cannot be captured in a single overlay model). The BP and BC moieties are labeled. The approximate positions of the pore entrance and selectivity filter relative to the overlaid blockers are shown for reference (red and blue horizontal lines, respectively).

The calculated solvation field within the blocker-accessible region of P is predicted to consist principally of H-bond depleted solvation (Figure 2), in agreement with our previously reported WaterMap results [10] (consistent with the largely non-polar lining of P). We note, however, that considerable rearrangement of the pore occurred during the unrestrained simulation (possibly resulting from truncation of the cytoplasmic domain, a question that will be addressed in future work).

**Figure 2.**
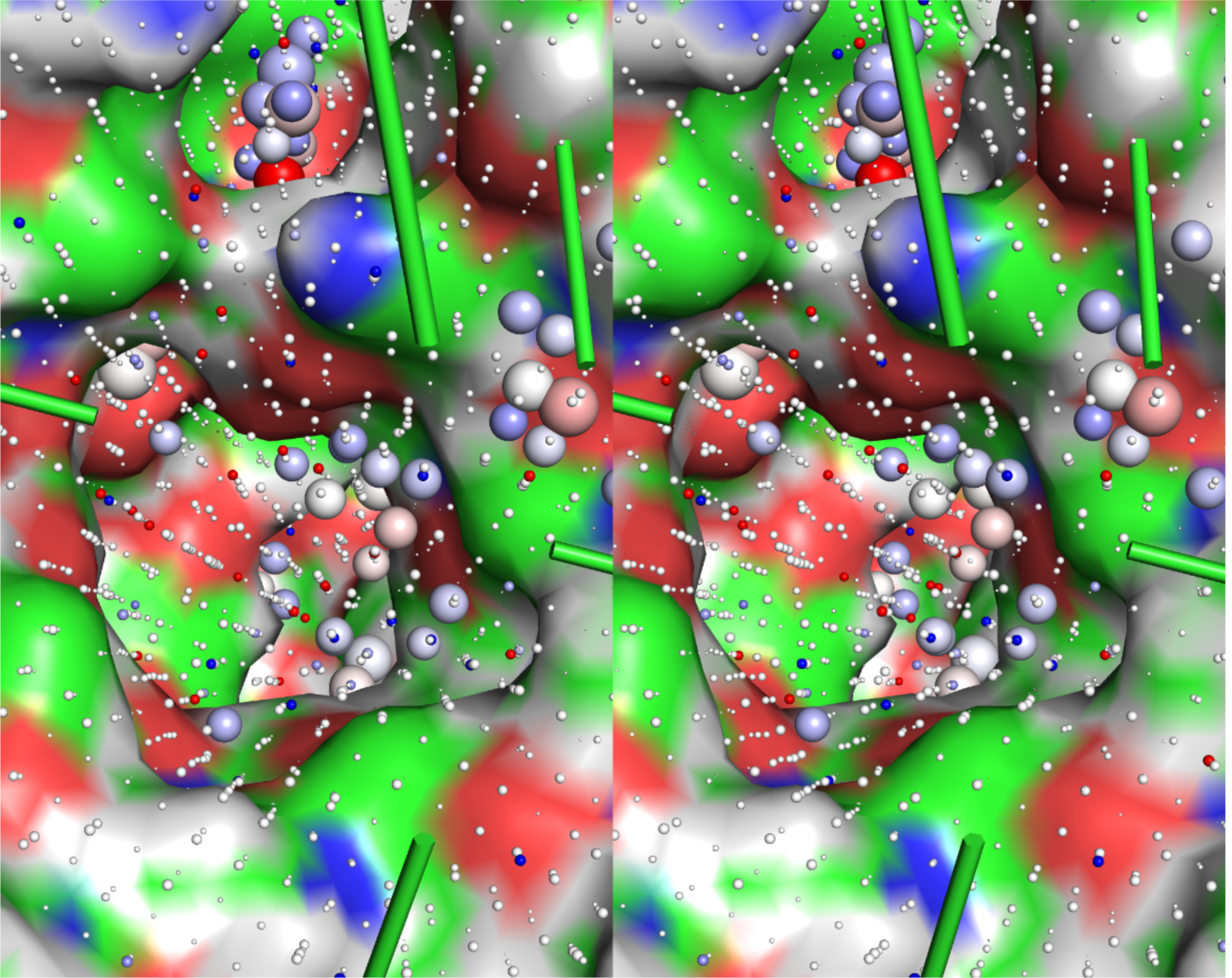
Cutaway stereo view of the solvation field within the pore of the time-averaged structure of the hERG channel (5VA1 with the cytoplasmic domain truncated), noting the significant rearrangement of the hERG structure during the unrestrained 100 ns SD simulation due in whole or part to the absence of the cytoplasmic domain. The solvation field consists primarily of ULOVs (consistent with the highly <u>non-polar</u> environment of P), accompanied by a small number of HOVs corresponding to weakly H-bond enriched solvation (the latter of which may be exaggerated by the loss of 4-fold structural symmetry of P during the simulation).

The solvation fields of the blocker conformations and charge state assumed in our overlay model (Figure 1) are shown in Figure 3, ordered by Redfern Class. All blockers are solvated by H-bond depleted water (reflected in numerous diffusely distributed ULOVs) that govern k_off_ from therapeutic targets and off-targets and k_-b_ from P. The relatively non-polar BP groups of the Class 1 and 2 blockers almokalant (Figure 3A), ibutilide (Figure 3B), and terfenadine (Figures 3F) are largely devoid of HOVs, consistent with the low expected desolvation cost of their BP moieties. The HOVs proximal to the aromatic groups in terfenadine and other blockers likely result from slight electrostatic attraction between water molecules and planar aryl groups unobstructed by tetrahedral hydrogens, the sizes of which are likely exaggerated in the absence of H-bonding. Conversely, the BP moiety of the Class 2 blocker terodiline (Figure 3G) is dominated by large HOVs surrounding the basic group, the desolvation cost of which seems more comparable to that of the Class 3 blocker flecainide (Figure 3H) and the Class 4 blocker desipramine (Figure 3K)) than other Class 2 blockers. This inconsistency may be partially explained by the additive bradycardic and hERG blocking effects of this drug [11,27]. The t-butyl acid and hydroxymethyl groups of the BP moiety in fexofenadine (the primary metabolite of terfenadine) are spanned by numerous HOVs (Figure 3I), consistent with higher BP desolvation cost and the Class 4 designation of this drug. The basic group and hydroxymethyl groups positioned midway along the BP moiety of propafenone (Figure 3J) are likewise consistent with higher desolvation costs and the Class 4 designation of this drug. The predicted solvation field of verapamil (Figure 3M) is consistent with that of Class 2, rather than Class 5 blockers, which may be explained by the comparatively low maximum reported C_max_, together with compensatory blockade of the inwardly conducting Ca_v_1.2 channel (the therapeutic target of this drug).

**Figure 3.**
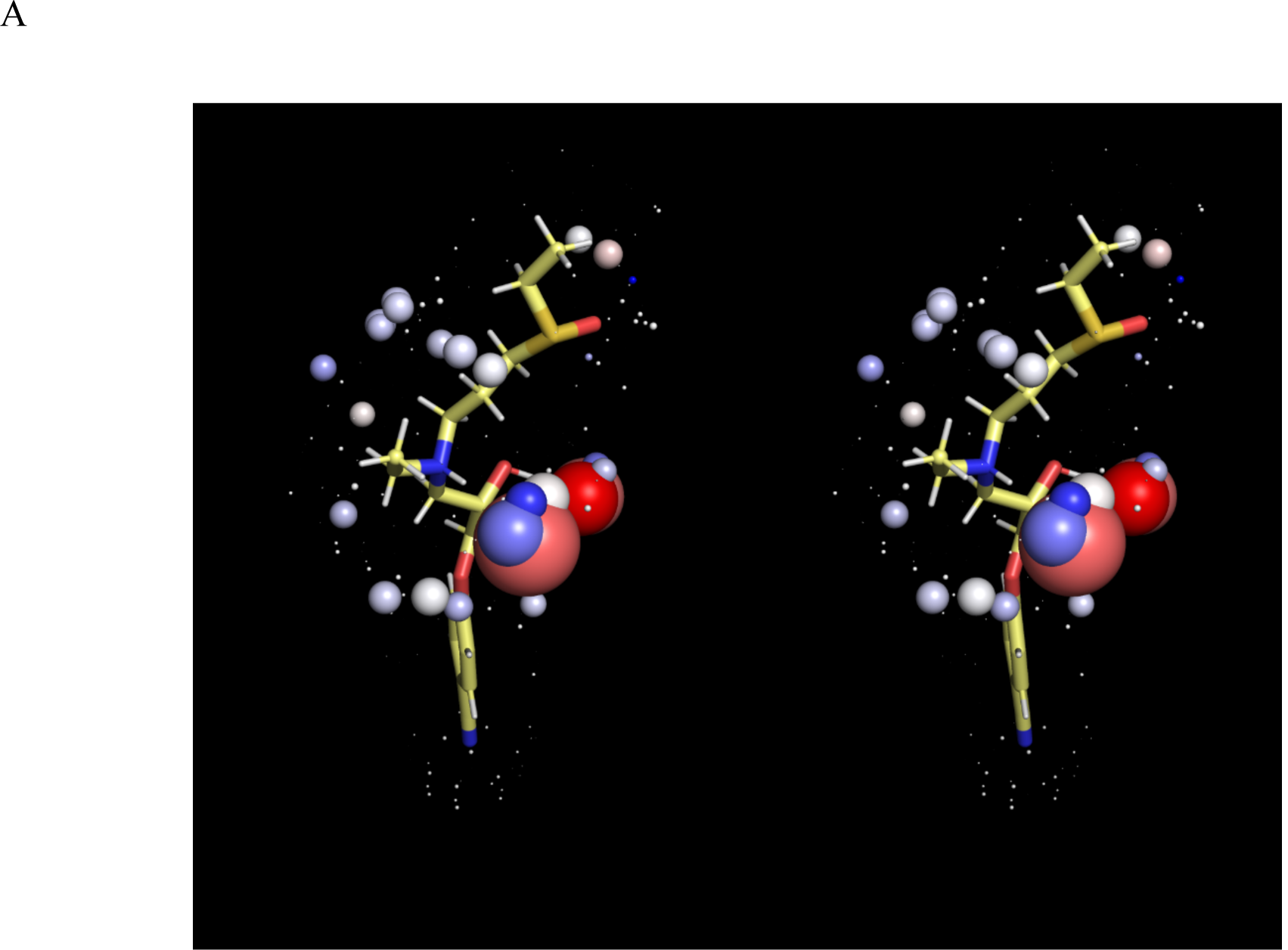
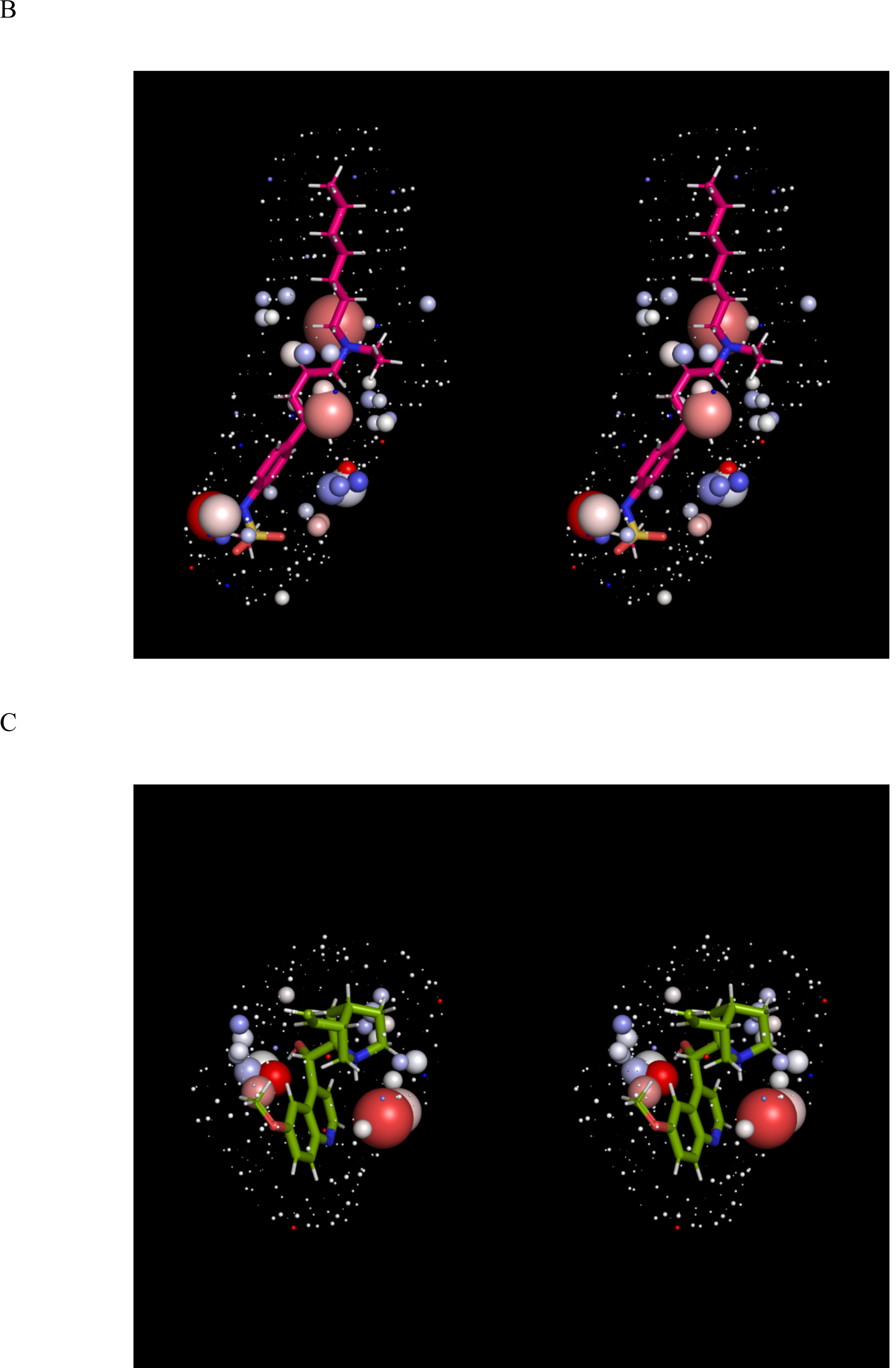
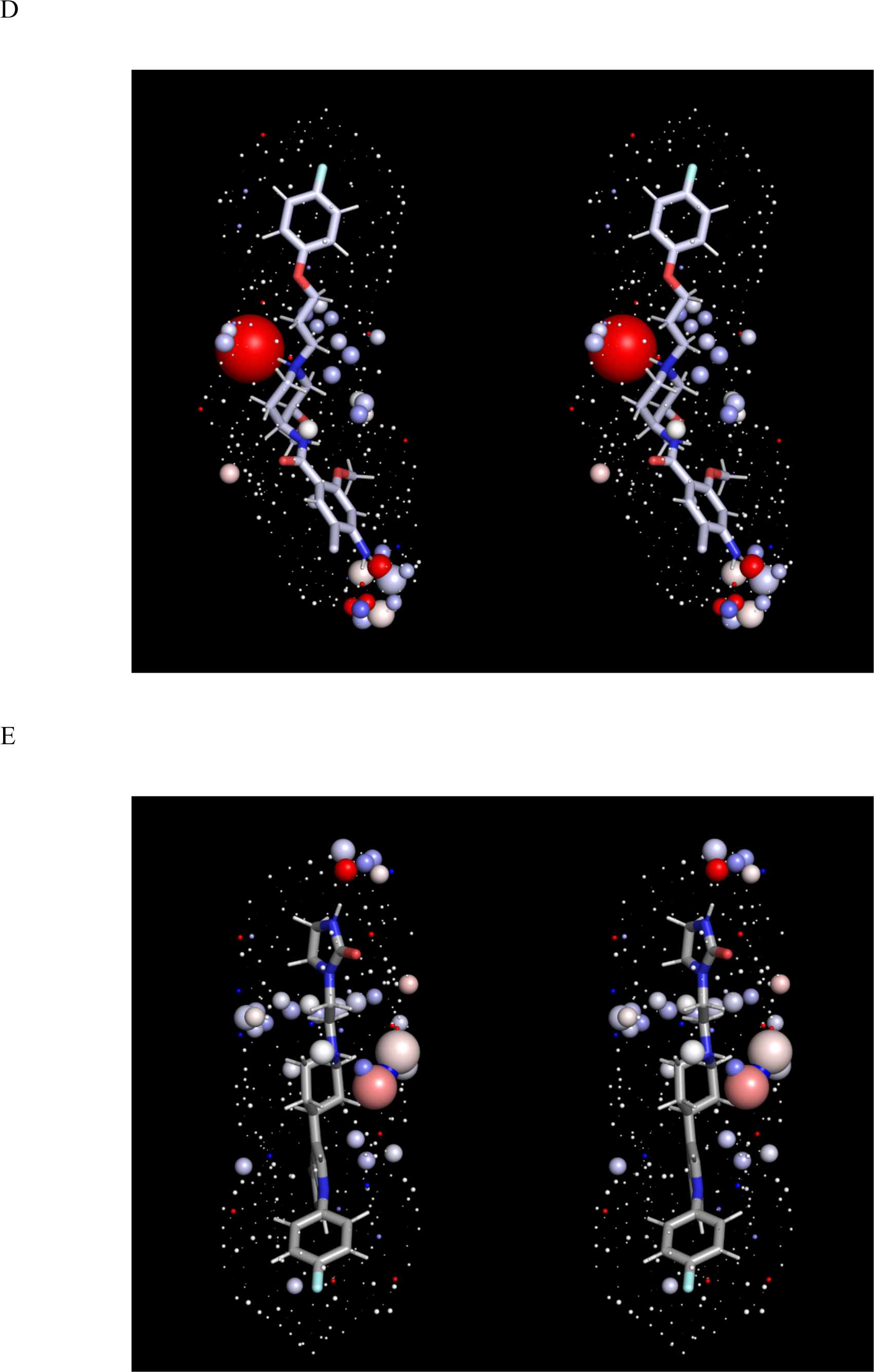
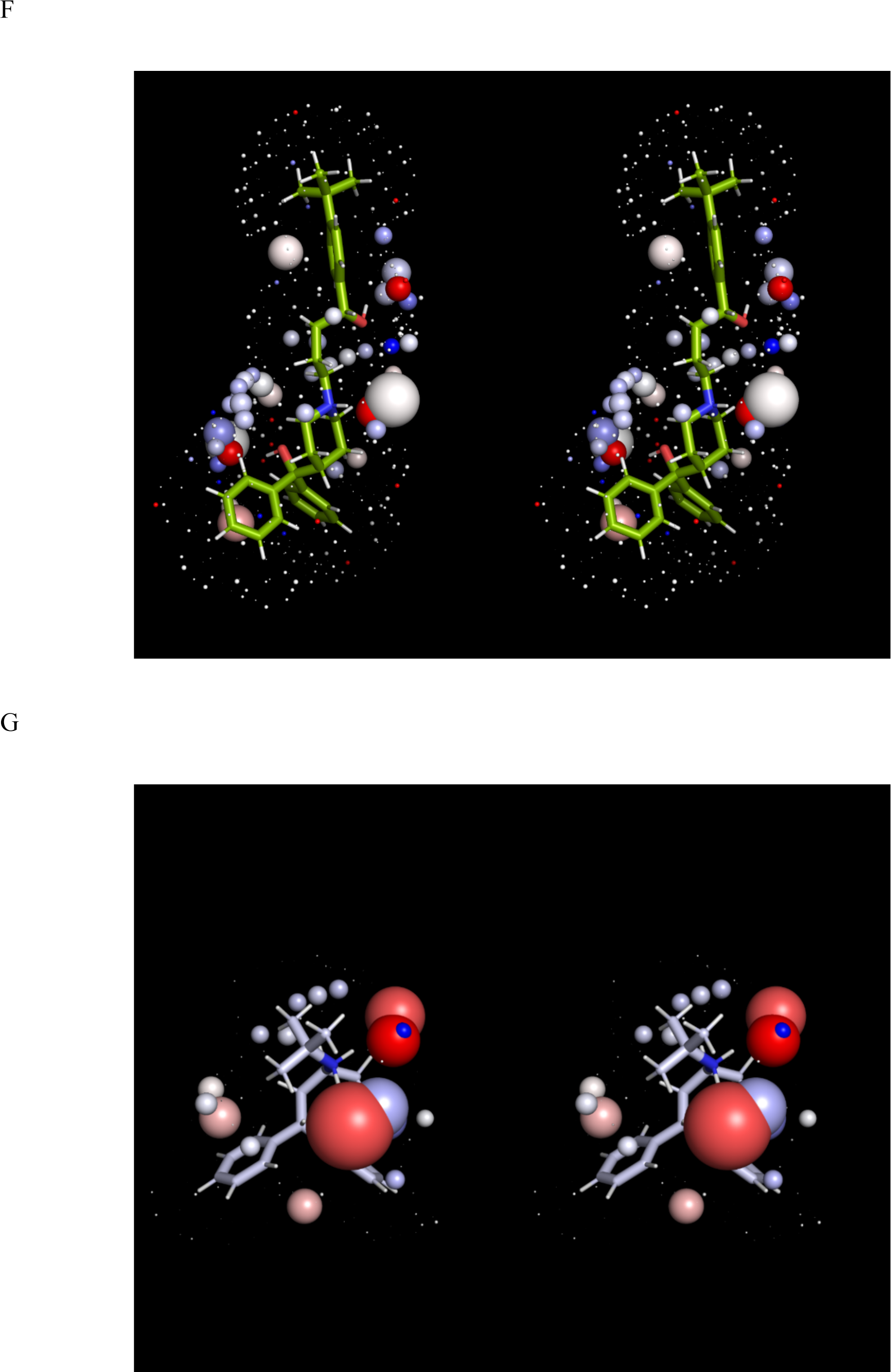
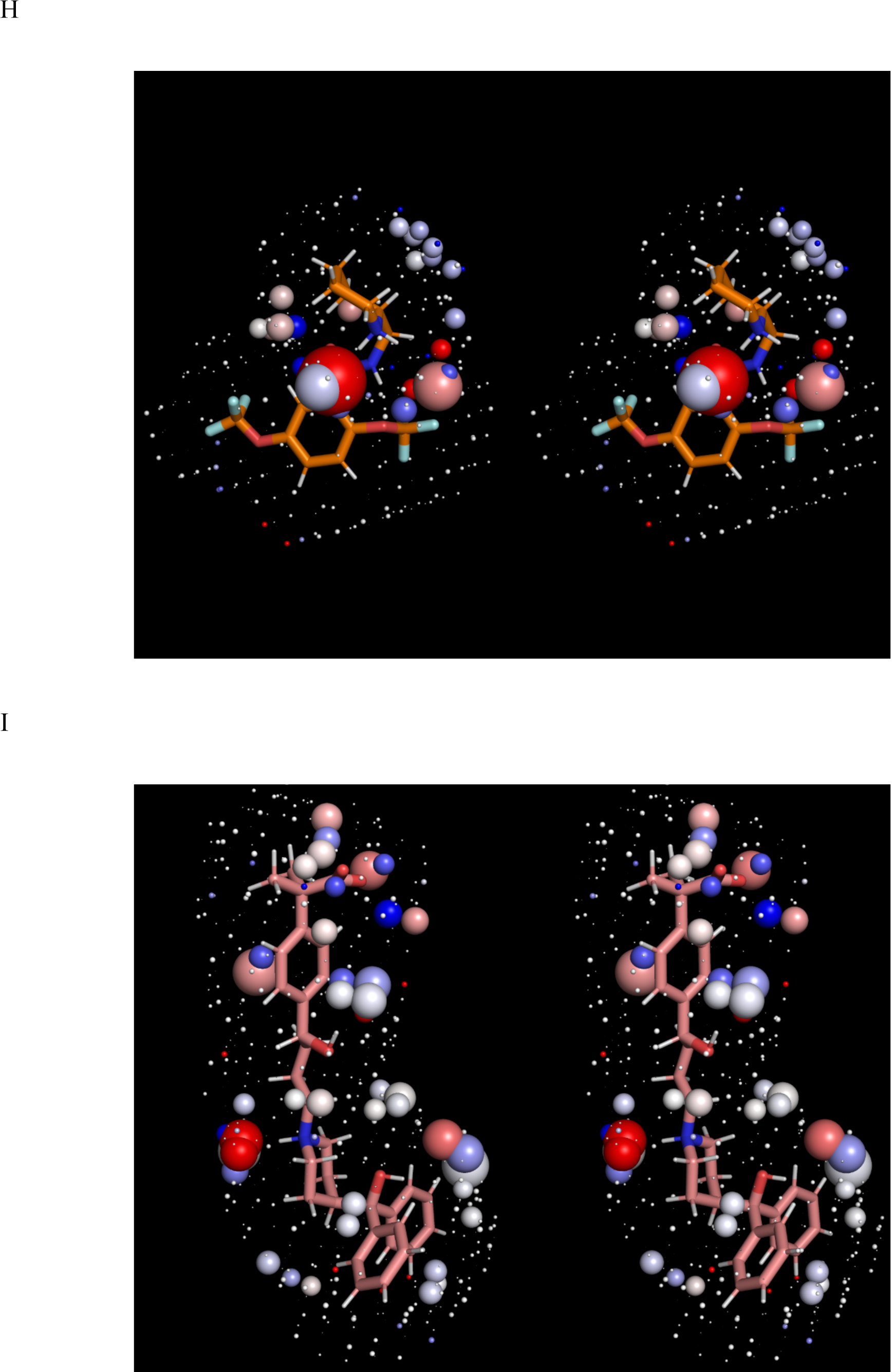
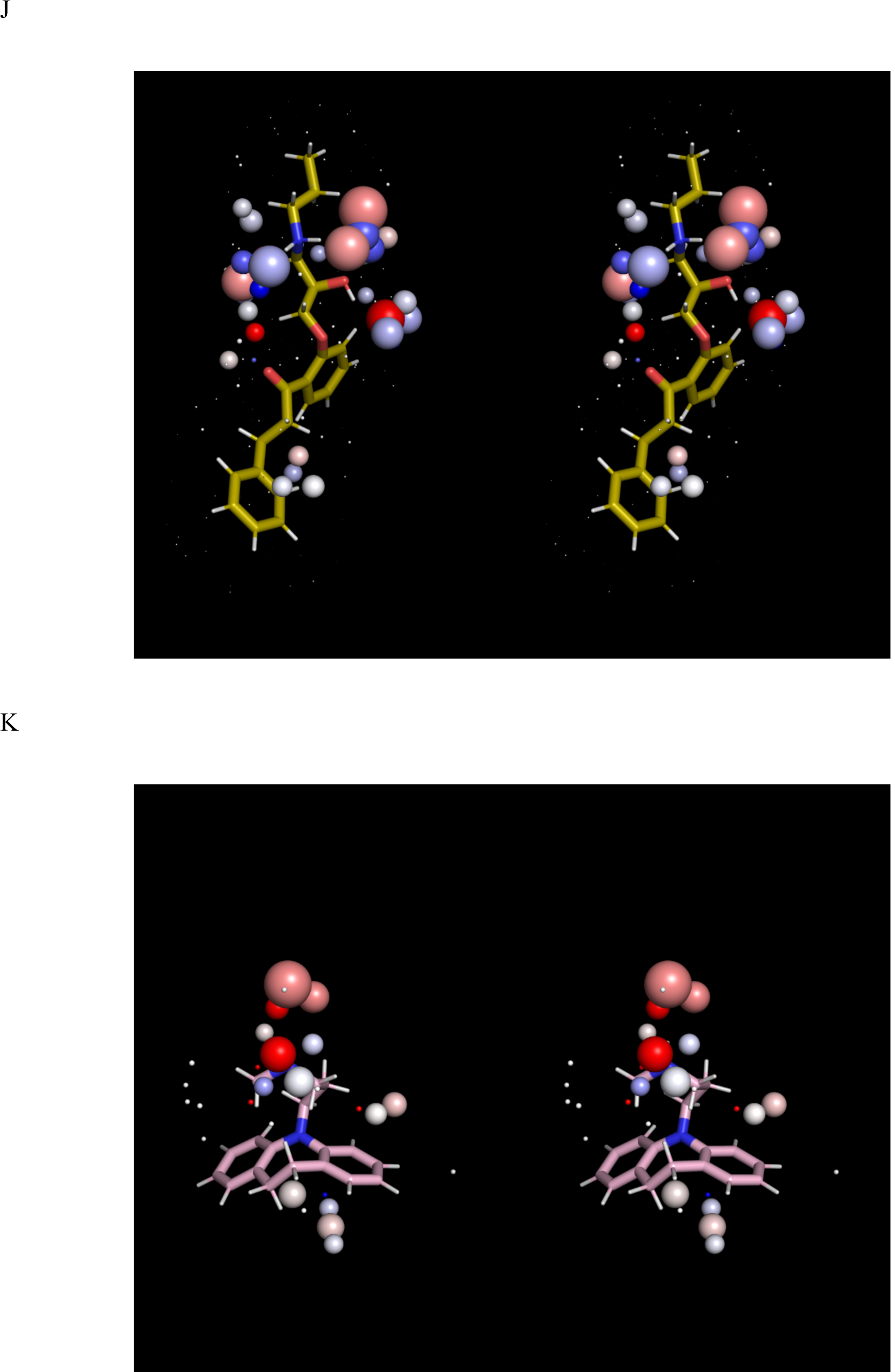
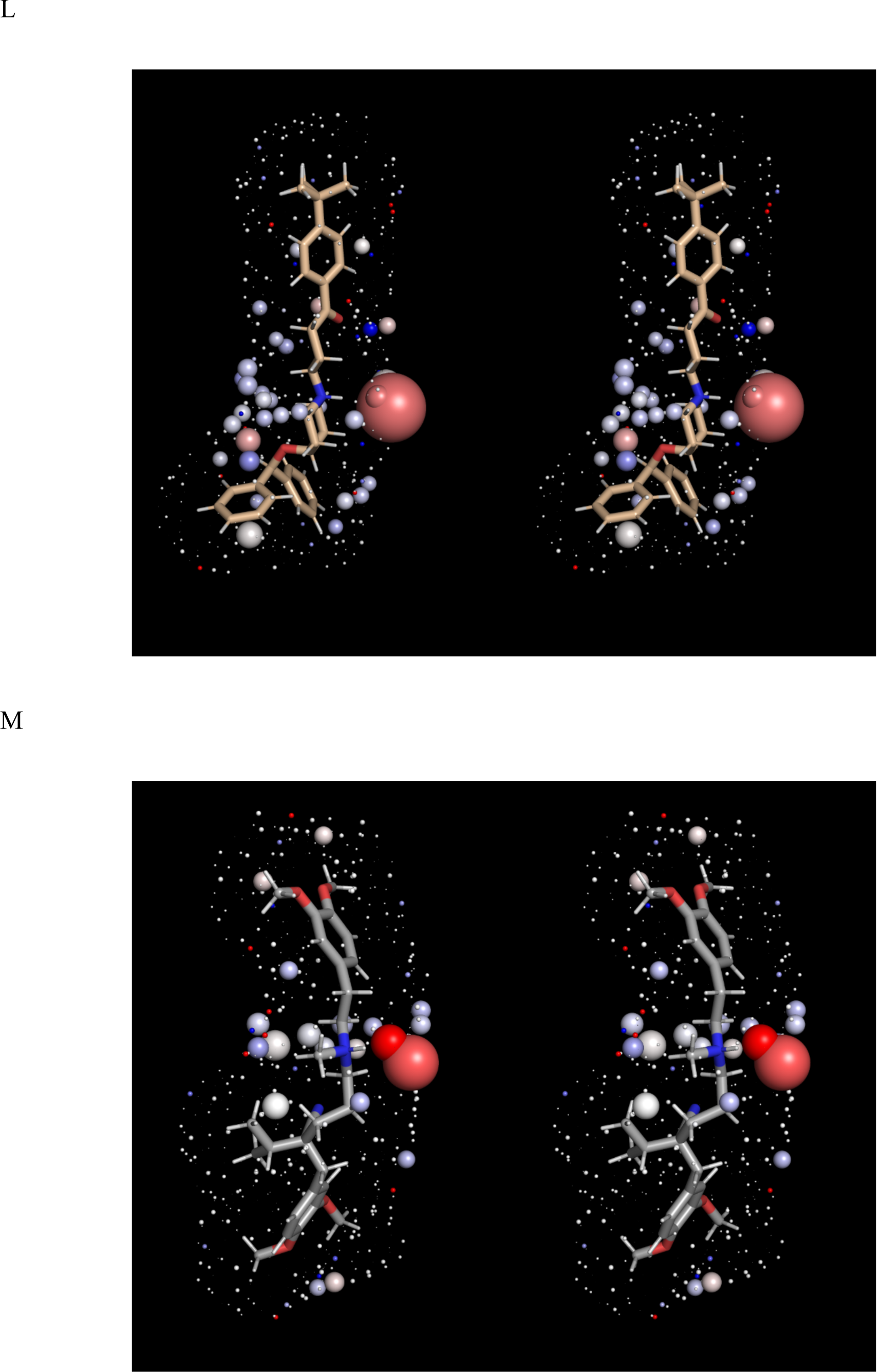
Stereo views of the solvation fields (represented by spheres as described in Materials and methods) and structures of the hERG blockers listed in Table 1 (sticks), the conformations of which correspond to those in Figure 1. The solvation fields of Class 1-2 blockers are highly similar and equate to low desolvation costs, with the exception of basic groups, which contribute heavily to solubility (as reflected in their larger HOV sizes). (A) Almokalant (Class 1). (B) Ibutilide (Class 1). (C) Quinidine (Class 1). (D) Cisapride (Class 2). (E) Sertindole (Class 2). (F) Terfenadine (Class 2). (G) Terodiline (Class 2). (H) Flecainide (Class 3). (I) Fexofenadine (Class 4). (J) Propafenone (Class 4). (K) Desipramine (Class 4). (L) Ebastine (Class 5). (M) Verapamil (Class 5).

The highly similar solvation fields of ebastine (Figure 3L) and terfenadine (Figure 3F) are inconsistent with their respective Class 5 and 2 designations, which is attributable to the large conformational difference between the t-butylphenylketone of ebastine (which is sterically incompatible with P) and the t-butylphenylmethane of terfenadine (Figure 4).

**Figure 4.**
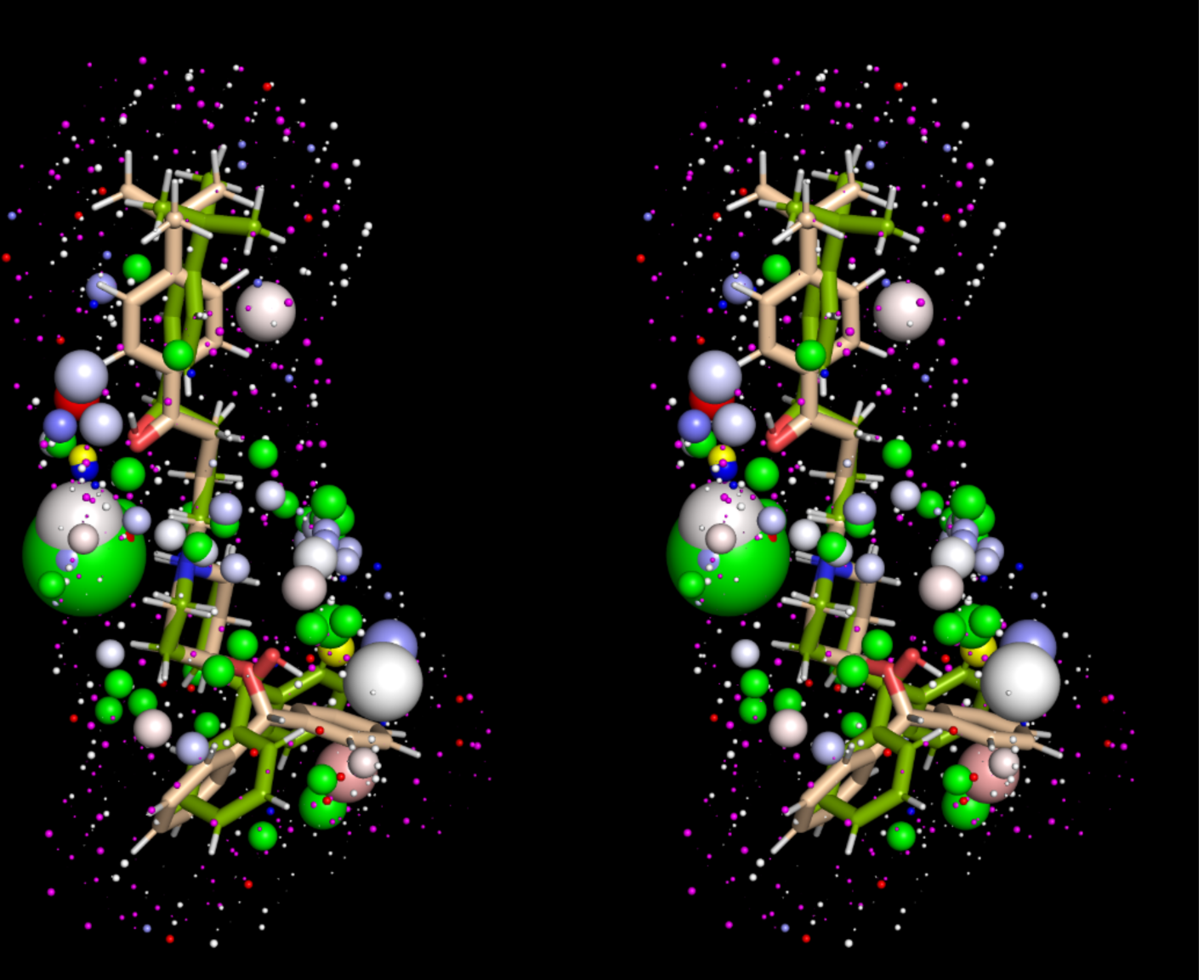
Stereo view of the solvation fields of ebastine (salmon sticks, green spheres = O preferred HOVs, yellow spheres = H preferred HOVs, magenta spheres = ULOVs) and terfenadine (green sticks, white spheres = ULOVs, standard red/white/blue spectrum = HOVs). The two blockers exhibit similar solvation fields despite their vastly different Redfern Class designations (5 and 2, respectively), leaving the significant conformational difference between the t-butylphenylketone of ebastine versus the t-butylphenylmethane of terfenadine to explain the large difference in hERG effects.

In addition, we tested the sensitivity of our results to the choice of conformation and charge state (Figure 5).

**Figure 5.**
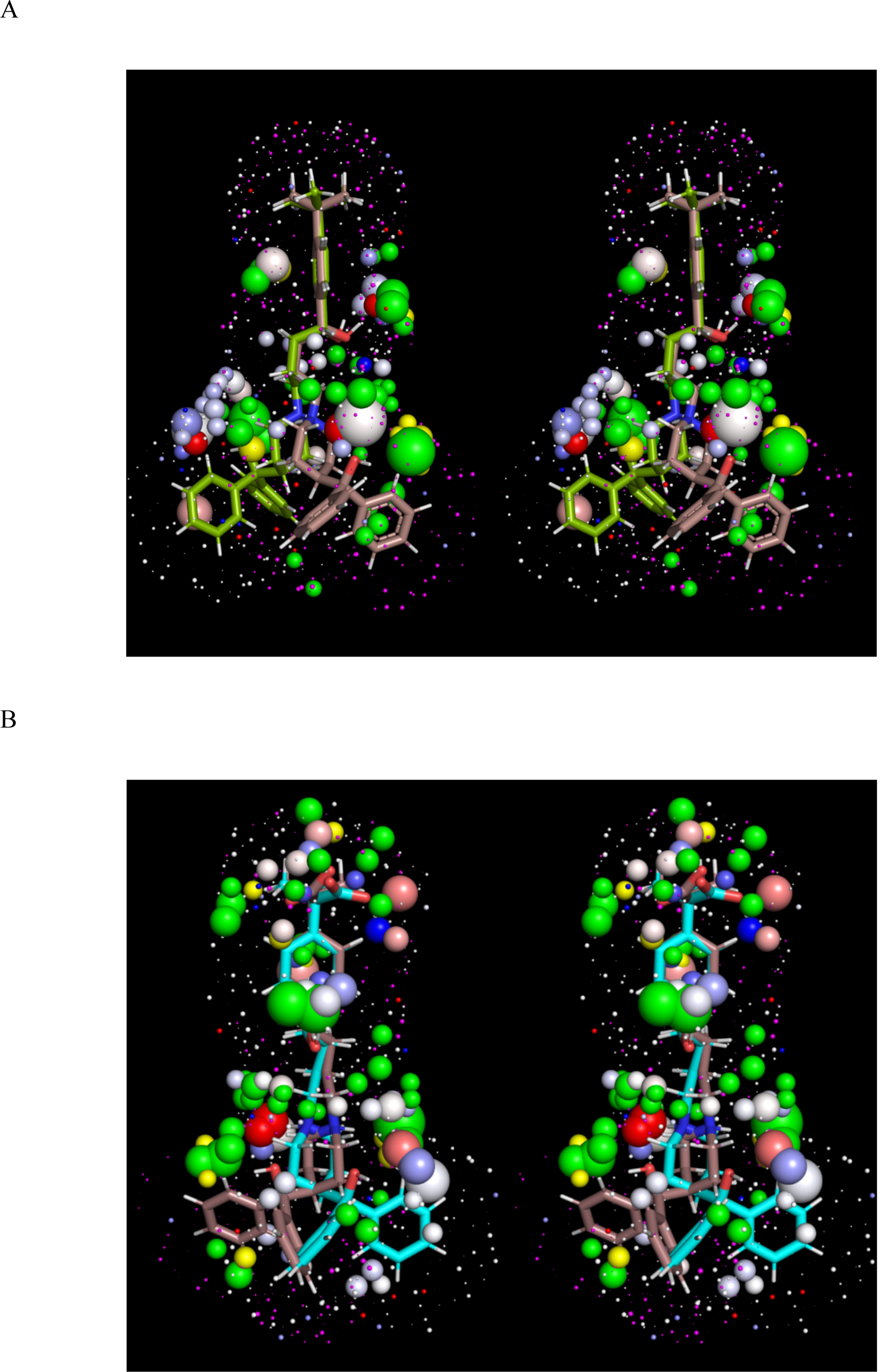
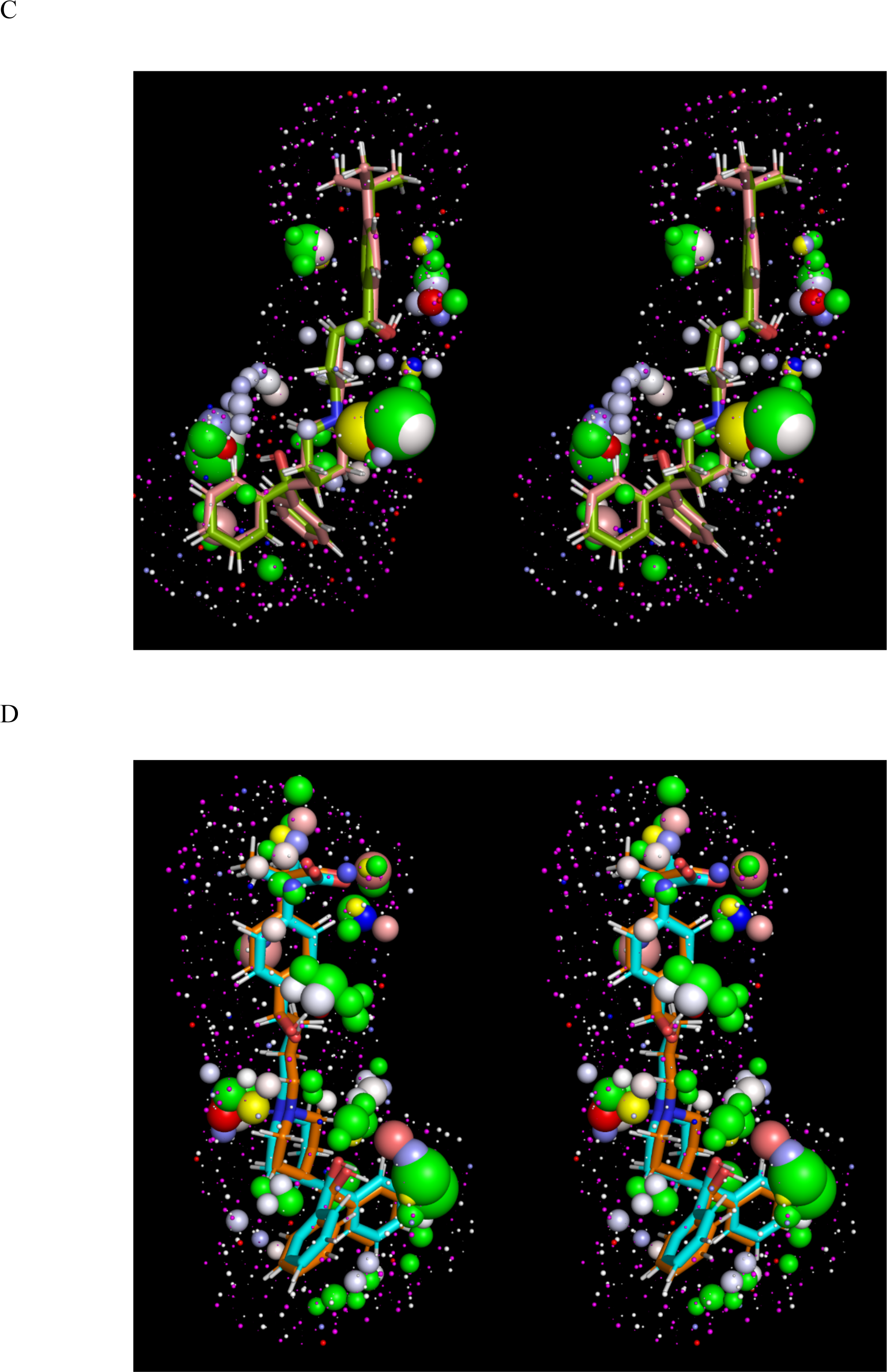
Stereo views of the solvation fields of alternate (though similar) conformations and neutral forms of selected blockers from Table 1 used to test the sensitivity of our results to conformational choice. (A) An arbitrary alternate terfenadine conformation overlaid on the conformation shown in Figure 1 (alternate form = salmon sticks, green spheres = O preferred HOVs, yellow spheres = H preferred HOVs, magenta spheres = ULOVs; reference form = green sticks, white spheres = ULOVs, standard red/white/blue spectrum = HOVs). (B) Same as A, except for fexofenadine (reference form = cyan sticks). (C) Neutral form of terfenadine overlaid on the charged form (neutral form = salmon sticks, green spheres = O preferred HOVs, yellow spheres = H preferred HOVs, magenta spheres = ULOVs; charged form = green sticks, white spheres = ULOVs, standard red/white/blue spectrum = HOVs). (D) Same as C, except for fexofenadine (neutral form = orange sticks, green spheres = O preferred HOVs, yellow spheres = H preferred HOVs, magenta spheres = ULOVs; charged form = cyan sticks, white spheres = ULOVs, standard red/white/blue spectrum = HOVs.

Overall, our WATMD results are consistent with our proposed straddled BP-in/BC-out binding paradigm, and in general agreement with the Redfern classification (with the explainable exceptions highlighted above).

## Discussion

### hERG blockers undergo atypical binding

In our previous work [2], we proposed that most hERG blockers conform to a canonical Y-shaped scaffold (and subsets thereof), in which the stem and cap of the Y straddle the pore entrance between the antechamber and pore, respectively. We modeled this motif and the binding mode thereof based on the observed bound state of the Y-shaped detergent GDN in the cryo-EM structure of Na_v_1.4 (PDB code = 6AGF) [28]. However, this hypothesis is contradicted by subsequent experimental evidence and previous computational predictions that blockers are <u>fully buried</u> within the ion conduction pathway, including:

1. Our own earlier induced-fit docking results [10,29].
2. The MD simulations of Dickson et al. [13], in which the authors predicted that both blockers and activators are fully bound within the pore.
3. Trapping of MK-499 within closed hERG channels inferred by Mitcheson et al. based on the observed slow recovery from block [30].
4. The first cryo-EM hERG structures solved by Wang et al. [24], in which the authors proposed that blockers occupy some or all of the “hydrophobic pockets” residing proximal to the intracellular side of the selectivity filter.
5. A recent cryo-EM structure of astemizole-bound hERG solved by Asai et al. [12] (7CN1).
6. Recent cryo-EM structures of flecainide and quinidine-bound Na_v_1.5 (6UZ0 and 6LQA, respectively), in which both blockers are fully buried within the pore (as claimed for astemizole-bound hERG and may be assumed for other hERG blockers as well) [25,31].

It is nevertheless reasonable to question whether modeled and experimentally determined structures solved <u>under equilibrium conditions</u> at high blocker concentrations recapitulate the <u>physiologically relevant</u> blocker-bound states of these channels, in which blocker-driven equilibration is likely precluded by the dynamic forces on the S5 and S6 helices by the voltage-sensing domain and membrane dipole potential [32]. It is likewise reasonable to question whether the structural states of bound blockers under native conditions are differentiable on the basis of electrophysiology data alone. Both fully and partially buried trappable blocker scenarios can exhibit extended recovery times since, unlike for non-trappable blockers, k_-b_ is not usurped by channel closing (which governs dissociation exclusively from the open state). **It is apparent that under fully native conditions, channel gating imposes severe time constraints on blocker association and dissociation that may be many fold weaker under in vitro conditions**. Blocker association and dissociation under <u>physiological</u> conditions are limited to:

1. The open/activated/inactivated states of hERG, which span the ∼350 ms duration of the cardiac AP (noting that the pore is likely accessible in the inactivated state of hERG).
2. The decaying open/activated sub-population of Na_v_1.5 channels that conducts the late Na^+^ current throughout the AP duration (noting that the peak current conducted by the fully populated open state of Na_v_1.5 is limited to the initial ∼2 ms of the AP and that the pore is likely inaccessible in the inactivated state).

The <u>fully buried</u> blocker-bound states observed in the cryo-EM and modeled structures are inconsistent with the aforementioned timescales for the following reasons:

1. Steric constraints on the passage of the bulky, chemically diverse BC moiety of most blockers through the pore entrance, which can be reasonably assumed to depend on time-consuming induced-fit rearrangements. The pore entrance is partially closed in the apo Na_v_1.5 structure (Figure 6A), and quinidine and flecainide are trapped behind the closed activation gate in their respective structures (Figures 6B and C, respectively). The fact that blocker association is limited to the open state serves as further evidence for equilibrated channel populations in the cryo-EM preparations.
2. Time-consuming induced-fit rearrangements between quinidine, flecainide, and the backbone/side chains of the Na_v_1.5 blocker binding site (assessed by comparison of 6LQA and 6UZ0 with the apo 6UZ3 (Figures 7A and B, respectively)). Relatively minor astemizole-driven induced-fit is apparent from a comparison of apo (5VA1) and astemizole-bound hERG (7CN1) structures (Figure 7C). However, the authors omitted the blocker from the PDB file for unexplained reasons, and we could not recapitulate the bound state described in reference 12 by docking the compound into the original 7CN1 structure using Glide XP (see Materials and methods).

**Figure 6.**
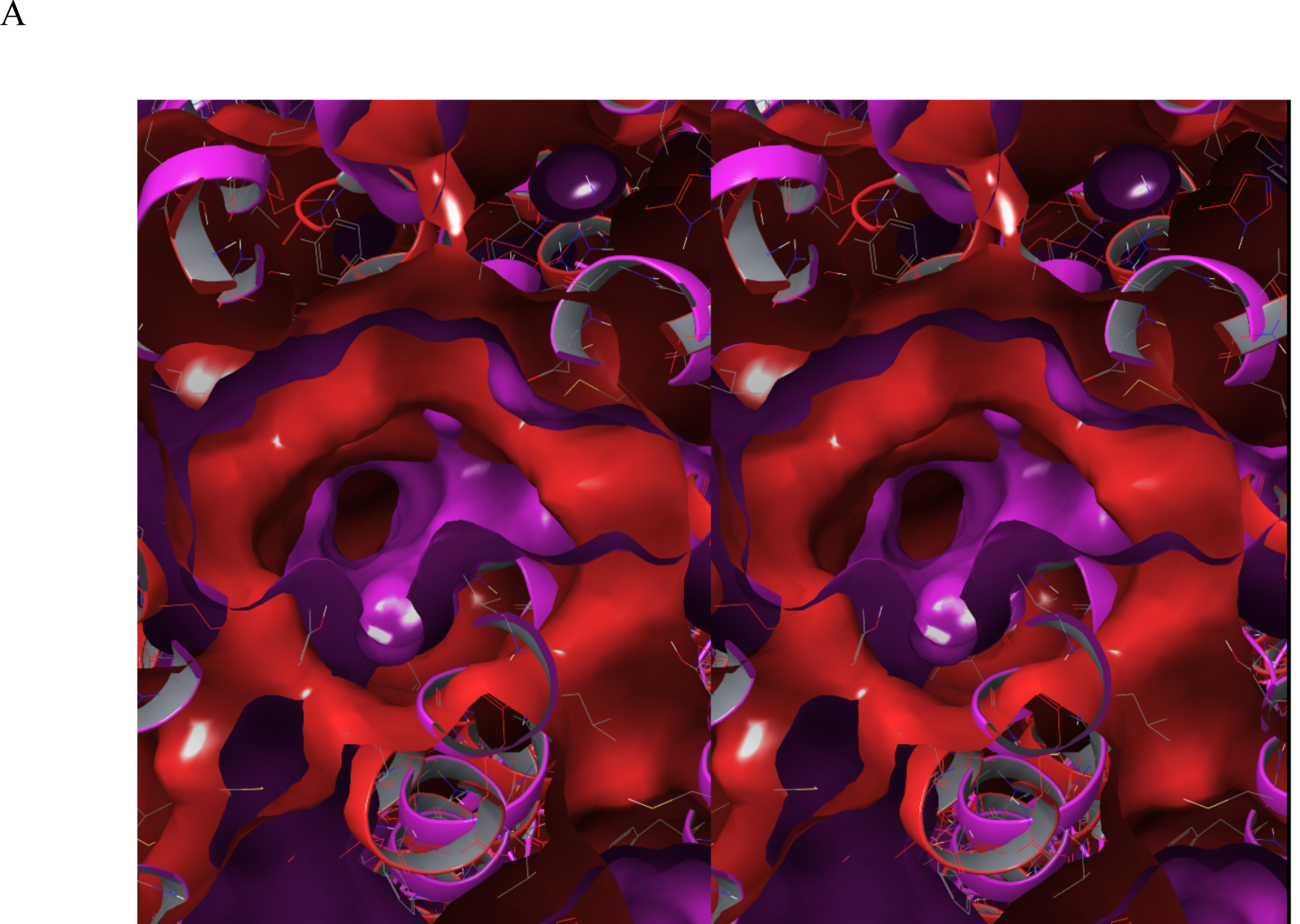
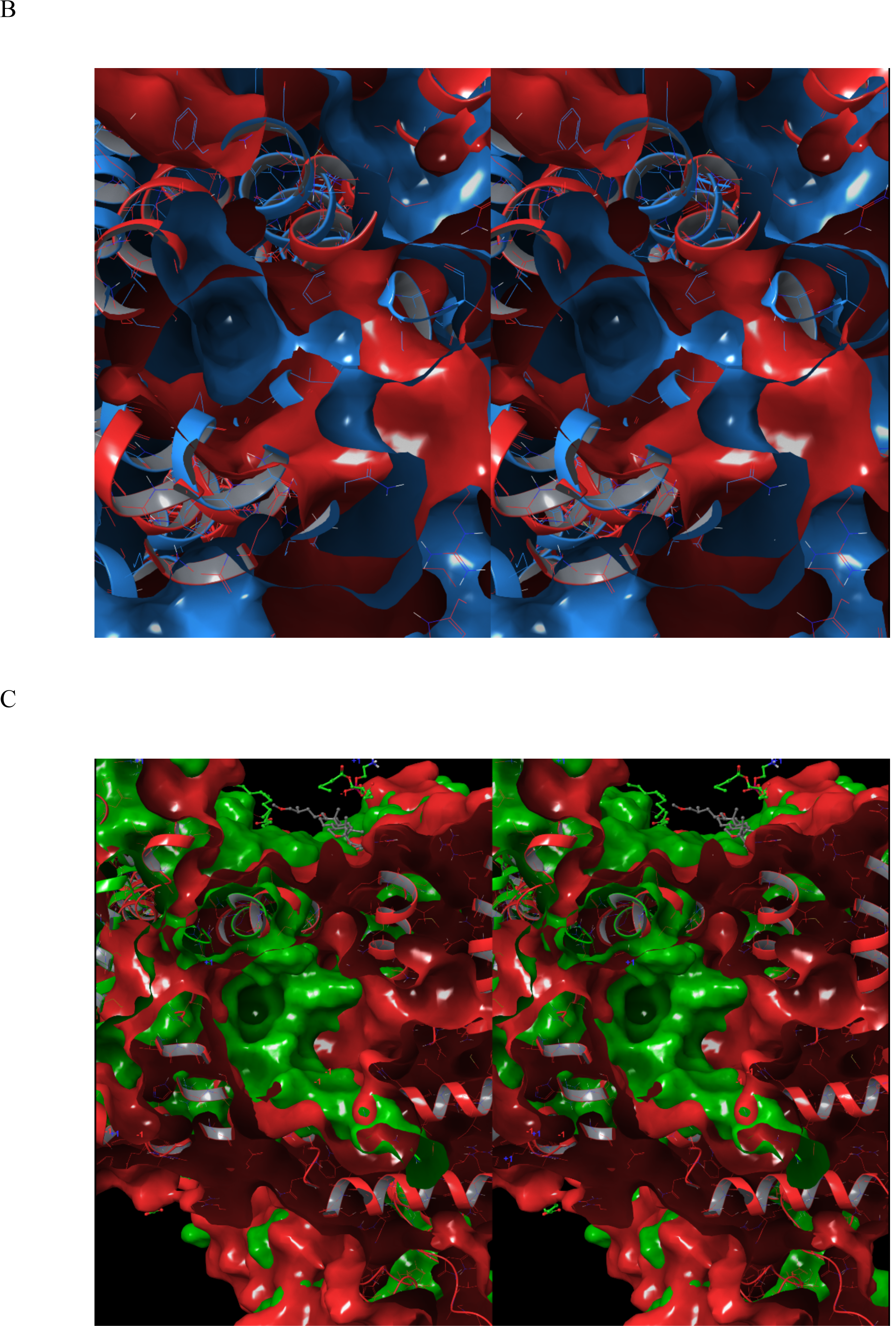
Stereo views comparing the pores of apo hERG with apo and bound Na_v_1.5 cryo-EM structures, looking from the cytoplasmic to extracellular direction. (A) The open state of apo hERG (5VA1, red) overlaid on the partially closed state of apo Na_v_1.5 (6UZ3, magenta). (B) Same as A, except for quinidine-bound Na_v_1.5 (6LQA). (C). Same as A, except for flecainide-bound Na_v_1.5 (6UZ0). The pore entrance in apo Na_v_1.5 is a small fraction of that in hERG, and nearly fully closed in the bound forms (consistent with equilibration).

**Figure 7.**
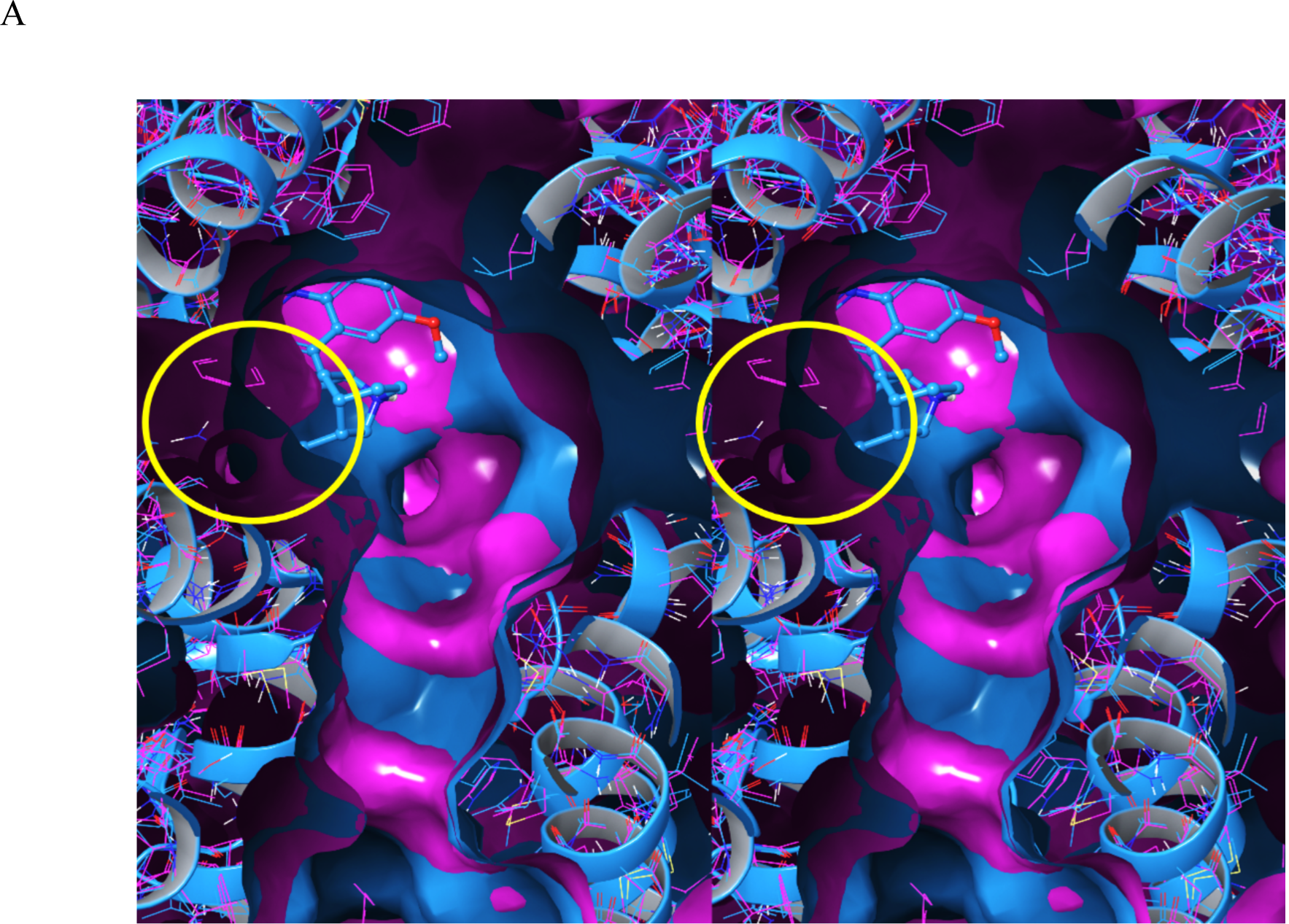
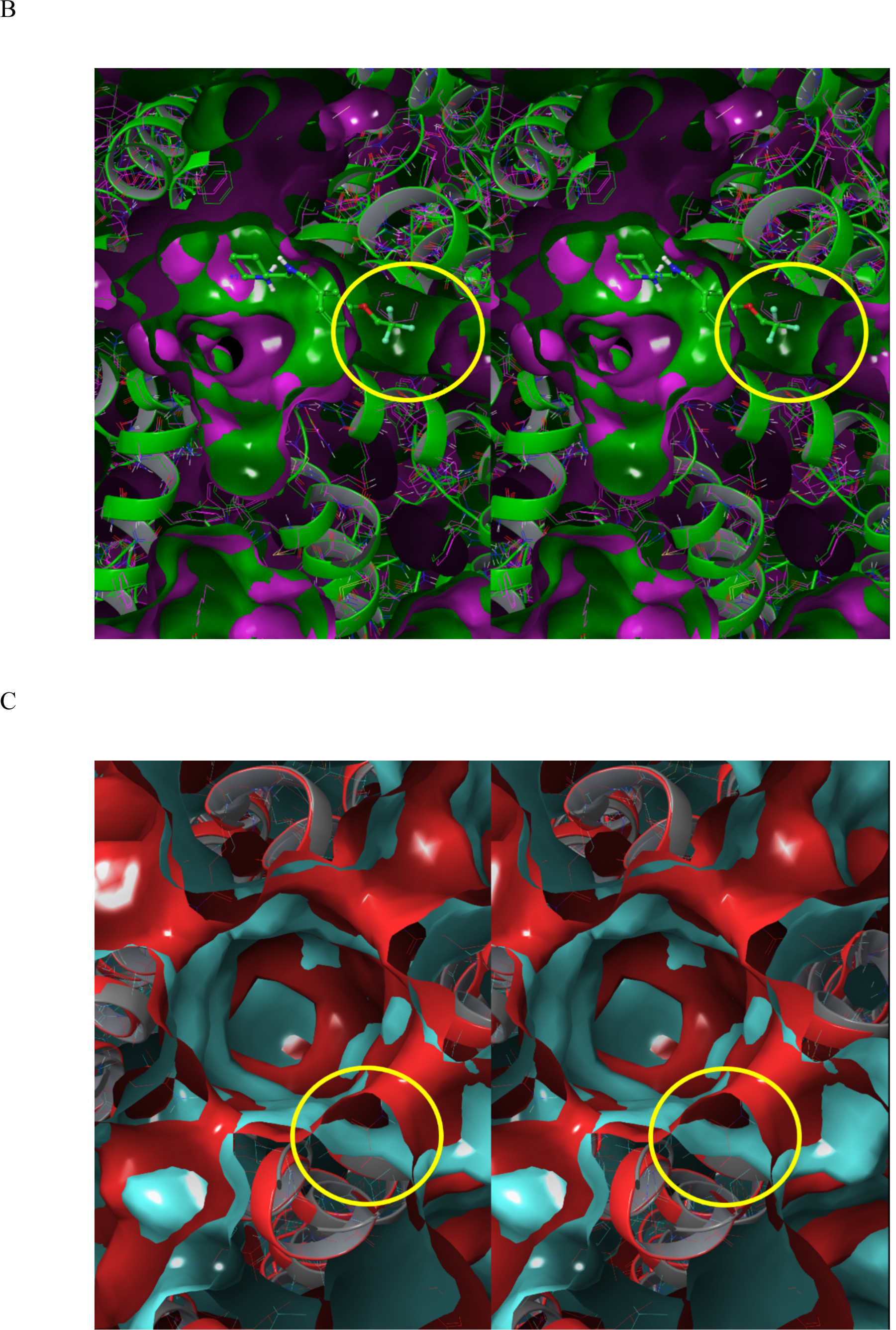
Stereo views of the pore in Na_v_1.5 (6UZ3/apo, 6LQA/quinidine, and 6UZ0/flecainide) and hERG (5VA1/apo and 7CN1/astemizole), looking from the cytoplasmic to extracellular direction (induced-fit rearrangements are circled in yellow). (A) Quinidine-bound Na_v_1.5 overlaid on apo Na_v_1.5. (B) Flecainide-bound Na_v_1.5 overlaid on apo Na_v_1.5. (C) Astemizole-bound hERG overlaid on apo hERG (noting the absence of astemizole in the structure). The primary difference is limited largely to the lower right region, which may or may not result from induced-fit.

### The <u>fully buried</u> blocker-bound states observed in the cryo-EM structures are additionally

questioned by the following discrepancies:

1. The lack of an obvious explanation for trappable versus non-trappable blocker subtypes based on the observed binding modes in 7CN1, 6LQA, and 6UZ0 compared with our proposed straddled BP-in/BC-out binding paradigm [2]. Time-consuming induced-fit dependent association and dissociation steps seem particularly implausible for non-trappable blockers, the occupancy of which necessarily builds and decays within each AP cycle (noting that trappable blockers likewise associate and dissociate dynamically during the open channel time window, rather than being irreversibly bound).
2. The lack of an obvious desolvation path from the fully occupied pore and one or more of the hydrophobic pockets in hERG. The pore volume is only partially filled by blocker BP moieties in our model (analogous to a syringe in which the plunger diameter < barrel diameter) (Figure 8A). We postulate that water expelled during blocker association flows through the unoccupied pore volume between the blocker surface and pore lumen anti-parallel to the BP association direction, exiting into the antechamber via the pore entrance (Figure 8B).
3. Overweighted interatomic contacts between P and BC in modeled structures based on force-field energies relative to the far greater solvation free energy contribution in our solvation free energy model.

**Figure 8.**
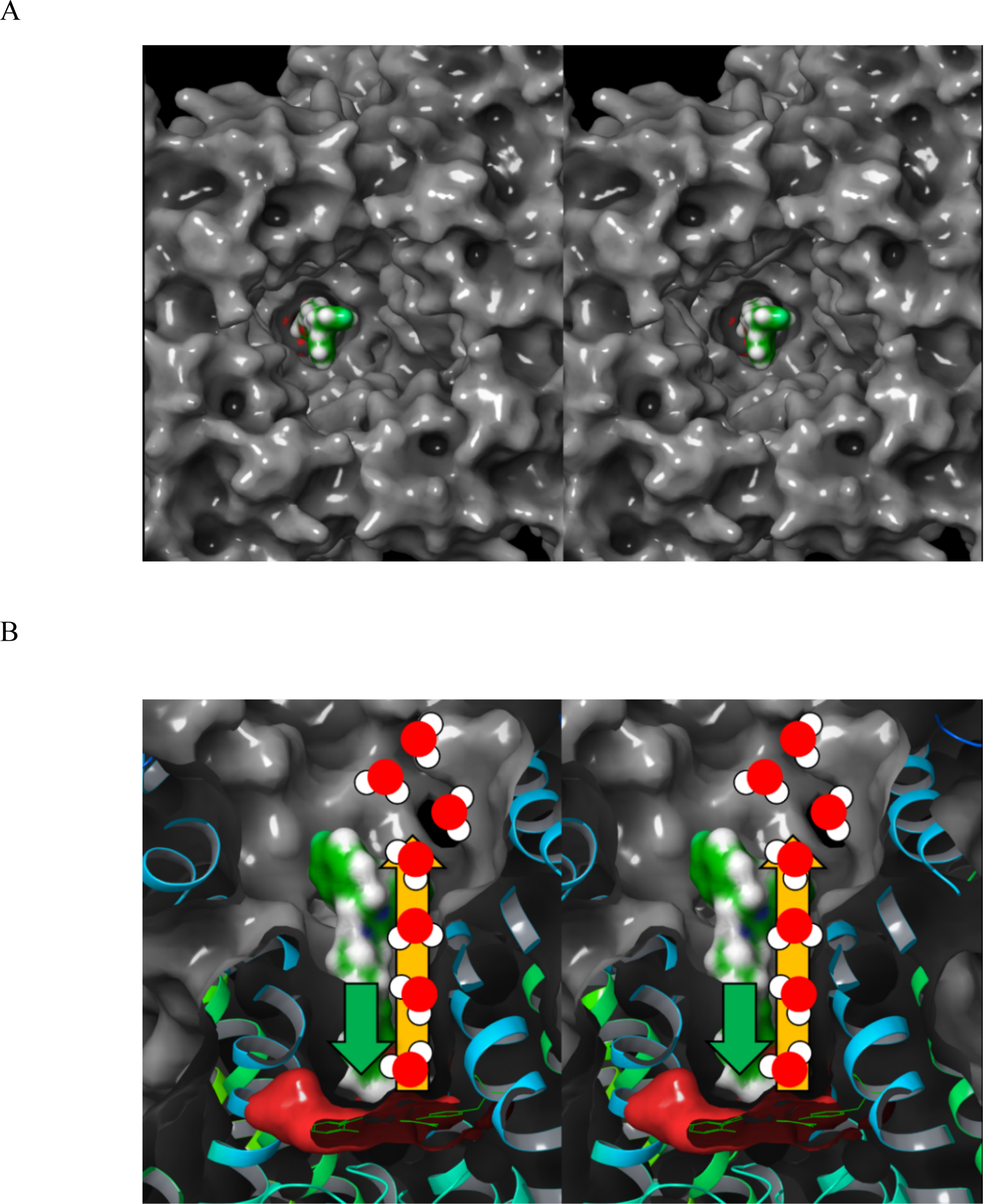
(A) Stereo view of the free pore volume between the surface of manually docked astemizole (green surface) and the pore surface (gray surface) viewed from the antechamber entrance looking along the pore axis toward the selectivity filter located deep within the cavity. This volume serves as the putative desolvation path between the leading edge of the associating blocker and antechamber. (B) Same as A, except as a cutaway viewed perpendicular to the pore axis, with the entrance at the top and selectivity filter at the bottom of the figure. The putative desolvation path is annotated by the orange arrow pointing from the leading edge of astemizole to the antechamber. The blocker association direction is annotated by the green arrow.

Instead, we propose that the buried blocker binding geometries observed in the hERG and Na_v_1.5 cryo-EM structures are relevant to <u>equilibrium</u> radioligand binding (RLB) assays and docked models (the latter of which are biased toward maximal interatomic contacts and minimum force-field energies), but not <u>non-equilibrium</u> conditions in vivo or in patch clamp assays performed at the native gating frequency. We showed previously that %hERG inhibition is poorly correlated among 7,231 compounds tested in RLB and QPatch assays, consistent with a variable, condition-dependent blocker binding paradigm (the observed/modeled fully buried versus partially buried states assumed in our model). Furthermore, the need for specifically positioned polar blocker groups capable of replacing the H-bonds of H-bond enriched protein solvation is obviated by the lack of HOVs within the pore (corresponding to “gatekeeper” solvation). As such, multiple binding geometries of a given blocker are conceivable under both equilibrium and non-equilibrium conditions. Our results are consistent with the high tolerance of hERG for diverse blocker chemotypes conforming to a general Y-shaped motif or any subset thereof. In the absence of H-bond enriched pore solvation, blocker association free energy costs are relegated to the <u>BP</u> desolvation cost, which is non-attenuated in the absence of H-bond replacements by the pore lining. No specific conformational or pharmacophore requirement exists for blocker binding. **As such, hERG blockade represents a form of non-specific binding** that depends principally on:

1. Steric shape/size pore complementarity (including the energetic preference for complementary BP and P conformations).
2. The total BP desolvation cost (including the availability of a desolvation path between the pore and bulk solvent).
3. Solubility (the gain in free energy between the solvated and un-solvated forms, which is typically enhanced by incorporation of one or more basic groups in BP).
4. Permeability (blocker desolvation and resolvation costs vis-à-vis the membrane desolvation cost).

### General hERG safety criteria suggested by our findings

The holistic optimization of primary target activity, permeability, solubility, and mitigation of hERG and other off-target activities, depends on the correct understanding of blocker structure-kinetic and structure-free energy relationships leading to the arrhythmic tipping point of hERG occupancy under native cellular conditions. Numerous studies aimed at predicting binding geometries and interactions between hERG and diverse blocker chemotypes via ligand- and structure-based methods have been attempted over the last few decades by ourselves and other workers [13,33]. Chemical mitigation nevertheless remains a largely trial-and-error proposition due to heavy reliance on:

1. Equilibrium binding and electrophysiology data that does not apply to binding sites undergoing high frequency buildup and decay cycles.
2. Status quo data-driven free energy models that are based on interatomic contacts (van der Waals, electrostatic, H-bonds, ρε-ρε, ρε-cation, hydrophobic, hydrophilic, etc.) rather than solvation fields (the putative horse’s mouth of SAR).

**The arrhythmic tipping point resides at a dynamic hERG occupancy of 50-60% (depending on the gating cycle length)** [10,11] **in simulations that we performed previously using a modified version of the O’Hara-Rudy model of the undiseased ventricular AP** [8,9]. Cellular arrhythmia in the form of atypical depolarizations can occur stochastically during transient or sustained occupancy incursions at or above this level. Occupancy amplitude increases in both cases as blocker exposure approaches the intracellular free C_max_, and decays with clearance-driven cytoplasmic decay. Measured IC_50_ weighted toward k_-b_ < the channel deactivation rate and k_b_ < the channel activation rate may result in overestimated potency of both trappable and non-trappable blockers. The arrhythmic occupancy by <u>trappable</u> blockers, which we take as ∼50% (the putative worst case scenario) depends on the blocker free C_max_ relative to the hERG IC_50_ (accumulating to the arrhythmic level at exposures ≈ the true blocker IC_50_ [10,11]), where the <u>rate</u> of fractional occupancy buildup depends on the slower of the antechamber and pore association steps. Trappable blocker dissociation is “paused” during the closed channel state, which exists during the approximately last two-thirds of the cardiac cycle (noting that decay of the bound state prior to channel deactivation results when k_-b_ exceeds the channel closing rate). Arrhythmic occupancy by <u>non-trappable</u> blockers, which builds and decays within each channel gating cycle, is governed by the following contributions:

1. Free intracellular blocker exposure ≥ hERG IC_50_ (concentration-driven) and/or k_b_ approaching the rate of channel activation (k_b_-driven), where k_b_ depends largely on the cost of expelling H-bond enriched blocker solvation <u>in the absence of polar pore replacements</u>. Free intracellular exposure, in turn, depends on:

a. Free plasma exposure, which depends on solubility, absorption, clearance, distribution, and dynamic plasma protein binding (PPB).
b. Cell permeability, which depends on plasma concentration and mutual membrane and blocker desolvation costs.
c. The fraction of blocker bound to membranes, lysosomes, and intracellular non-hERG off-targets.
2. k_-b_ approaching the rate of channel deactivation, which depends largely on the rate of channel closing or blocker/pore resolvation costs that are qualitatively proportional to the number of ULOVs surrounding P and BP (whichever is faster).

**Knowledge of blocker trappability is therefore essential for assessing concentration-occupancy relationships based on in vitro measurements. Furthermore, data models unwittingly generated using mixtures of trappable and non-trappable blockers may not be meaningful (noting that trappable and non-trappable analogs may occur within the same chemical series, as evidenced by the propafenone analogs reported by Windisch et al.** [7]).

Optimal hERG occupancy is achieved within a Goldilocks zone of solubility, permeability, and binding for both trappable and non-trappable blockers, as follows:

1. Solubility is proportional to the H-bond free energy of H-bond enriched solvation, as reflected in HOV number and size (noting that all HOVs fall deep within the tail of the Gaussian distribution of water counts tabulated by WATMD). Poor solubility reduces the free blocker concentration, whereas high solubility equates to high desolvation costs that hamper both drug-target and drug-off-target binding. H-bond enriched solvation is further enhanced by basic groups, which additionally speed k_b_ (as a function of increasing pKa) due to electrostatic attraction with the negative field in P. Since blocker potency is typically enhanced by basic groups, it follows that the electrostatic k_b_-speeding contribution of such groups typically outweighs the k_b_-slowing desolvation contribution.
2. Permeability is proportional to the desolvation cost of H-bond enriched blocker solvation, which depends additionally on polar groups for replacing the H-bond enriched solvation of membrane phospholipid head groups (the underlying mechanism reflected in the Pfizer Rule of 5). The on-rate to the antechamber is proportional to the <u>intracellular</u> blocker concentration. High desolvation costs corresponding to HOVs located anywhere on blocker surfaces slow permeation, as does the lack of polar groups that facilitate membrane desolvation (again, which is qualitatively consistent with the Rule of 5).
3. The overall asymmetric distribution of HOVs among BP and BC blocker moieties is consistent with our proposed binding mode, in which BP (exhibiting the lower desolvation cost/lower association free energy barrier) projects into the pore, while BC (exhibiting the higher desolvation cost/higher association free energy barrier) remains within the solvated antechamber. HOVs on the BP moiety of most blockers (except fexofenadine) are positioned around the basic group (when present), which is typically incorporated for solubility enhancement purposes, and in some cases, therapeutic target binding. The Redfern Classes are correlated qualitatively with the basic group and HOV position/desolvation cost along the longitudinal BP axis, as follows:

a. A lower desolvation cost is incurred when positioned near the proximal end of BP in blocker Classes 1-3 (corresponding to the pore entrance region).
b. A higher desolvation cost is incurred when positioned near the middle or distal end of BP in blocker Classes 4-5 (projecting deeper within the non-polar pore environment).

The Goldilocks zone of pro-arrhythmic hERG occupancy thus depends on optimal solubility + permeability + free intracellular exposure + fast antechamber k_on_ + fast pore k_b_, as reflected in the HOV sizes and distributions among the BP and BC regions. Our WATMD results are qualitatively consistent with the aforementioned dependencies. The Class 1-2 blockers in our dataset (almokalant, ibutilide, terfenadine) (Figures 3A, 3B, 3F) lack large HOVs on the BP moiety but contain significant numbers of ULOVs needed to slow k_-b_ (in addition to ULOVs distributed around the surface of P). Conversely, HOVs are predicted at the BP positions of Class 4 blockers (fexofenadine, propafenone, and desipramine) (Figures 3I-K). The t-butyl acid group located on the distal end of BP explains the weak hERG activity of fexofenadine, whereas the weaker potencies of propafenone and desipramine can be explained by the more distal position of the basic group on BP relative to that of Class 1-2 blockers. The HOV positions in all of these cases reside deeper along the pore axis, consistent with higher desolvation costs at these positions.

The data reported by Redfern et al. (Table 1) suggests that arrhythmic occupancy levels are achieved at the maximum end of the reported free C_max_ range, rather than the minimum end of the reported IC_50_ range for most of the Class 1-2 drugs in our study (with the exception of cisapride and terodiline) (Table 3 and Figure 9). Cisapride is a known non-trappable Class 2 blocker [6], the high pro-arrhythmicity of which is necessarily due to fast k_b_ and/or exposure escalation far above the therapeutic level. Since this drug does not achieve the putative ∼50% arrhythmic hERG occupancy threshold at the maximum reported IC_50_ and C_max_ (Tables 1 and 3), the former explanation is more likely (noting that fast k_b_ is consistent with the 2 nM lowest reported IC_50_). The Class 2 pro-arrhythmicity of terodiline (predicted to be trappable based on our overlay model [2]) likewise achieves the putative arrhythmic hERG occupancy level at the maximum reported C_max_ and minimum IC_50_ (Tables 1 and 3). Interestingly, verapamil exhibits sub-arrhythmic occupancies for all IC_50_-C_max_ combinations in Table 3, suggesting that the lack of pro-arrhythmicity of this drug is only partially attributable to concurrent Ca_v_1.2 blockade.

**Table 3.**
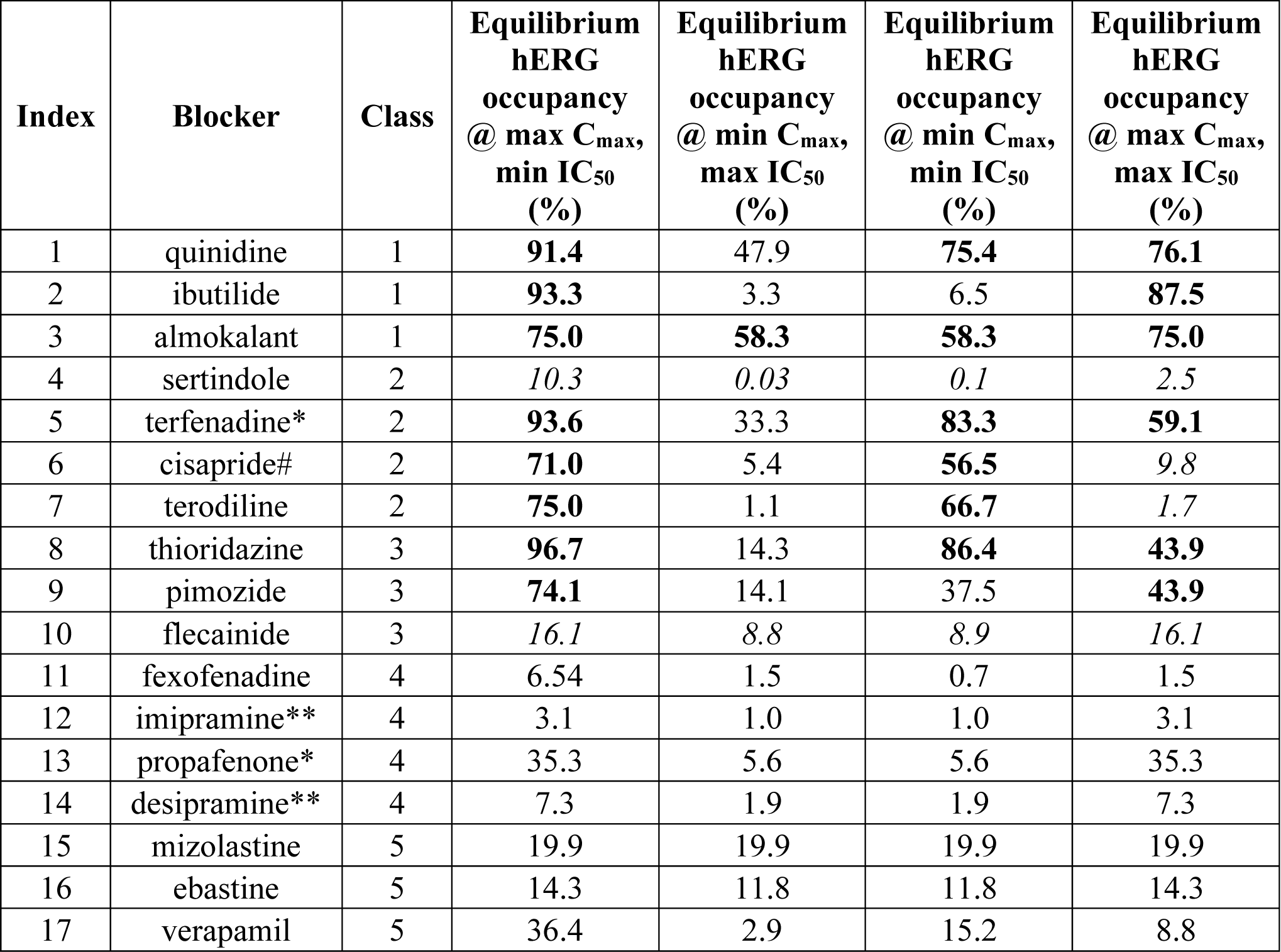
Equilibrium hERG occupancy (predicted from the Hill equation defined above) as a function of maximum and minimum reported C_max_ and hERG IC_50_. The Redfern Classes correlate poorly with reported max IC_50_ at min C_max_, and best with reported min IC_50_ at max C_max_. However, max IC_50_ and max C_max_ are also well-correlated with Redfern Class (bolded values), with the exception of cisapride and terodiline (italicized values), suggesting that arrhythmic occupancy is driven more by C_max_ escalation above the therapeutic level than by potency (noting that sertindole is an across the board outlier, which is also shown in italics).

**Figure 9.**
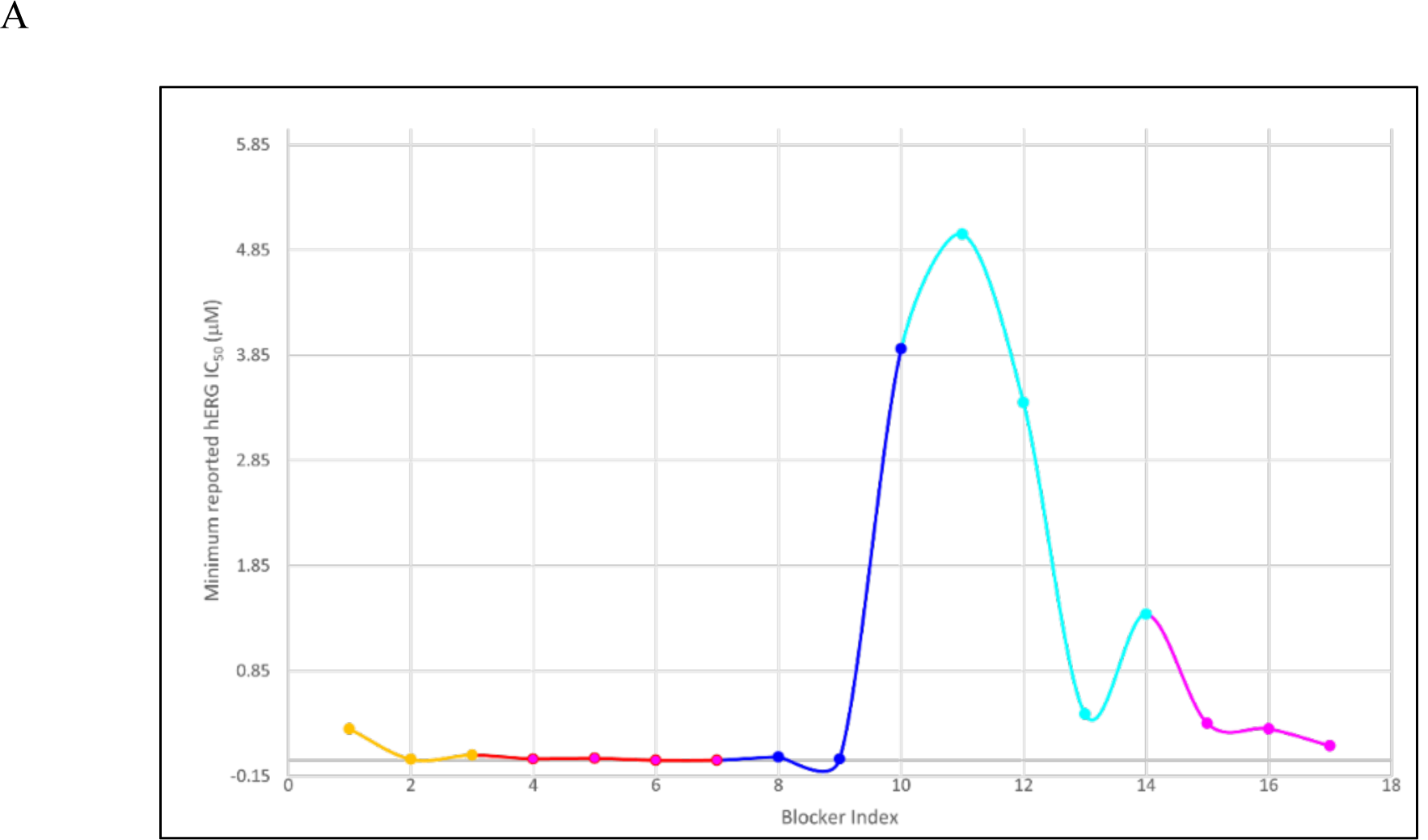
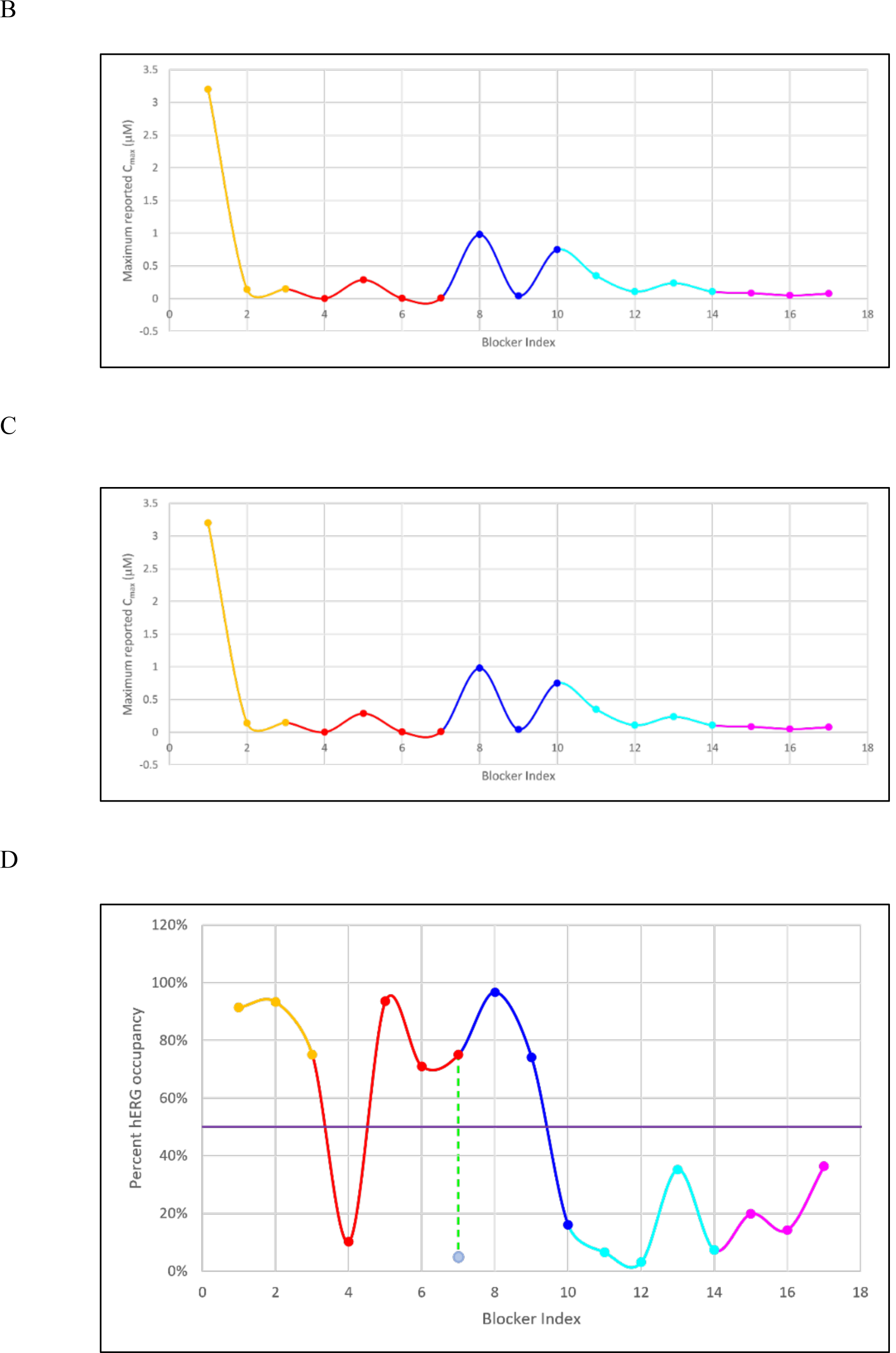
Parameter distributions listed in Table 1 annotated by Redfern class (orange = class 1, red = class 2, blue = class 3, cyan = class 4, and magenta = class 5). Mizolastine (omitted from our dataset) was added to increase the representation of class 5 blockers. (A) Distribution of the <u>minimum</u> reported hERG IC_50_ across the dataset (B) Distribution of the <u>maximum</u> reported C_max_ across the dataset. (C) Distribution of the ratios of the minimum reported hERG IC_50_/maximum reported C_max_ across the dataset. (D) Equilibrium hERG occupancy calculated using (max C_max_)/(max C_max_ + min IC_50_) (equation 1), with the 50% arrhythmic level predicted in our previous work [11] denoted by the purple line. The pro-arrhythmic effects of verapamil (blocker 17) are widely believed to be attenuated by co-blockade of Ca_v_1.2 channels. It is apparent from Table 3 and Figure 3M that verapamil approaches, but does not exceed, the putative arrhythmic ∼50% hERG occupancy threshold at the minimum reported IC_50_ and maximum reported C_max_, suggesting that co-blockade of the outward hERG and inward Ca_v_1.2 currents is only partially responsible for the safety of this drug. Sertindole is an outlier in the Redfern dataset, having reported incidences of arrhythmia, despite falling well below the ∼50% hERG occupancy at the maximum reported C_max_ and minimum reported hERG IC_50_. The calculated <u>equilibrium</u> hERG occupancy of cisapride (blocker 6) is likely exaggerated due to the known non-trappability of this compound. The calculated solvation field of terodiline is more similar to the Class 4 blockers desipramine and imipramine than Class 2 blockers, consistent with calculated occupancy = 1.7% at the highest reported IC_50_ = 0.7 μM (denoted by the green dotted line and light blue circle).

### The optimal hERG mitigation strategy suggested from our findings

Efficacious and toxic drug-target and drug-off-target occupancy are achieved at threshold free drug concentrations in the target and off-target compartments, which in turn, depend on threshold levels of solubility and permeability (both in terms of absorption in the case of oral drugs and permeability in the case of intracellular targets and off-targets). We have claimed in this and our other works that non-covalent binding, solubility, and permeability are all governed principally by solvation free energy, which in turn, is governed by the solvation fields of drugs, targets, off-targets, and membranes. The therapeutic index (TI) is defined as the ratio of the toxic to efficacious free drug concentration, where toxicity is due to occupancy of one or more off-targets like hERG. A safe TI is achieved when the separation between the efficacious and toxic free drug exposures is sufficiently wide to preclude adverse effects in humans (i.e., when the TI >> 1), allowing for unintended escalation of the free C_max_ above the therapeutic level due to drug-drug interactions (DDIs) or overdose (the circumstances under which all cases of reported arrhythmias occurred among the marketed drugs in the Redfern dataset). The objective of hERG mitigation is therefore to achieve a TI >> 1 by minimizing the efficacious and maximizing the toxic free drug exposure thresholds. We have claimed in this and our other works that non-covalent drug-target/off-target occupancy, solubility, and permeability are all governed principally by solvation free energy, which in turn, is governed by the solvation fields of the participating molecules. It then follows that lead optimization should be focused on achieving drug solvation fields residing at the intersection of the following (noting that target, off-target, and membrane solvation fields are fully determined by nature):

1. Maximum complementarity between the solvation fields of drugs and their targets at which efficacious occupancy is achieved.
2. Minimum complementarity between the solvation fields of drugs and all off-targets, such that toxic occupancy is avoided.
3. A threshold level of H-bond enriched solvation commensurate with a threshold solubility level.
4. A threshold balance between H-bond enriched and depleted solvation (reflected qualitatively in the Pfizer rule of 5) commensurate with a threshold permeability level through the gut (in the case of oral drugs) and cell membranes (in the case of intracellular targets).

In our previous work, we demonstrated that hERG binding occurs in a two-step fashion, consisting of a second order loading phase, in which blockers are captured in the antechamber (which does not depend on blocker desolvation due to the large volume of the cavity), followed by transient first order association of the blocker BP group and the open pore (which depends on transient desolvation and partial desolvation of the BP and BC groups, respectively) [2]. Our proposed canonical “BP-in/BC-out” blocker binding model, together with WATMD, can be used to predict the BP moiety of a given blocker, and <u>qualitatively</u> guide mitigation to hERG occupancies << the putative arrhythmic ∼50% threshold at the highest plausible free C_max_ by:

1. Achieving the highest <u>therapeutic target</u> occupancy at the lowest possible exposure (the biggest bang for the buck) via “kinetically tuned” drug-target binding [34]. **Low efficacious exposure affords the greatest possible safety margin for hERG and all off-target liabilities**, many of which may track with target activity (noting that second order antechamber loading depends on the intracellular drug concentration, whereas first order BP-P binding depends on k_b_ and k_-b_).
2. Disruption of blocker trappability (which effectively slows k_-b_) via the incorporation of a bulky group within the putative constriction zone in the closed channel state (as we reported previously [2]).
3. Disruption of P-BP shape compatibility (e.g., putatively exemplified by ebastine versus terfenadine).
4. Introducing strategically positioned polar groups on BP aimed at increasing the desolvation cost of this substituent and slowing k_b_ without slowing the primary target k_on_ (noting that increased polarity at other blocker positions, as reflected in decreased logP) is likely insufficient for hERG mitigation.
5. Exploring structure-solubility and permeability relationships, which are likewise governed by solvation free energy and membrane desolvation/resolvation costs.
6. Optimal positioning and pKa of a basic group on BP used to enhance solubility (noting that increased solubility as a f(pKa) results in higher desolvation cost, but also speeds k_b_ via favorable electrostatic interactions with the negative field residing within P).

## Conclusion

This work is a follow-on to our previous work in which we postulated that hERG blockers are first captured by the intracellular antechamber contained within the CNBH and C-linker domains of the channel, followed by the projection of a single R-group (BP) into the open pore (P) [2]. Blocker occupancy is powered principally by desolvation of the H-bond depleted solvation of P, BP, the pore-facing surface of BC, and the peri-pore region of the antechamber (which slows k_-b_). The key determinants of hERG blockade consist of:

1. Steric size/shape complementarity between BP and P, where BP is typically a non-polar/weakly polar rod-shaped moiety of almost any chemical composition. The antechamber volume is sufficient for capturing large chemotypes that need not fully fit within P.
2. Second order buildup of the antechamber-resident blocker population (governed by on-rate = k_c_ · [free intracellular blocker] · [free antechamber]), followed by first order buildup of the P-BP population (governed by k_b_ · [antechamber-bound blocker]).
3. A Goldilocks zone of BP solvation free energy needed to simultaneously achieve permeability, solubility, and low desolvation cost of P and BP during association (which applies to both the therapeutic target and all forms of off-target binding). P is predicted by both WaterMap and WATMD to contain H-bond depleted and bulk-like solvation, which incurs no desolvation cost during association, and a resolvation cost during dissociation equal to the total free energy of H-bond depleted solvation expelled during BP-P association.
4. A basic group residing somewhere within BP that is typically used to improve solubility (traded off against a higher desolvation cost). Electrostatic attraction between P and such groups speeds k_b_ more than the additional desolvation cost slows it.
5. Blocker trappability, in which arrhythmic hERG occupancy is more likely achievable by trappable blockers whose occupancy accumulates across multiple APs.

Successful hERG mitigation is necessarily achieved by slowing k_b_ via increased polarity and desolvation cost of BP (within the Goldilocks zone of solubility, permeability, and target binding), pKa attenuation that maintains solubility (and in some cases efficacious target occupancy), avoidance of trappability, and disruption of blocker-pore shape complementarity (e.g., ebastine).

